# Linked-read whole-genome sequencing resolves common and private structural variants in multiple myeloma

**DOI:** 10.1101/2021.12.09.471893

**Authors:** Lucía Peña-Pérez, Nicolai Frengen, Julia Hauenstein, Charlotte Gran, Charlotte Gustafsson, Jesper Eisfeldt, Marcin Kierczak, Fanny Taborsak-Lines, Remi-André Olsen, Ann Wallblom, Aleksandra Krstic, Philip Ewels, Anna Lindstrand, Robert Månsson

**Author notes:** **CONTACT INFORMATION:** Robert Månsson, Karolinska Institutet, Center for Hematology and Regenerative Medicine (HERM), Neo Building, floor 7, Blickagången 16, 141 52 Huddinge, Sweden.

## Abstract

Multiple myeloma (MM) is an incurable and aggressive plasma cell malignancy characterized by a complex karyotype with multiple structural variants (SVs) and copy number variations (CNVs). Linked-read whole-genome sequencing (lrWGS) allows for refined detection and reconstruction of SVs by providing long-range genetic information from standard short-read sequencing. This makes lrWGS an attractive solution for capturing the full genomic complexity of MM. Here we show that high-quality lrWGS data can be generated from low numbers of FACS sorted cells without DNA purification. Using this protocol, we analyzed FACS sorted MM cells from 37 MM patients with lrWGS. We found high concordance between lrWGS and FISH for the detection of recurrent translocations and CNVs. Outside of the regions investigated by FISH, we identified >150 additional SVs and CNVs across the cohort. Analysis of the lrWGS data allowed for resolving the structure of diverse SVs affecting the MYC and t(11;14) loci causing the duplication of genes and gene regulatory elements. In addition, we identified private SVs causing the dysregulation of genes recurrently involved in translocations with the IGH locus and show that these can alter the molecular classification of the MM. Overall, we conclude that lrWGS allows for the detection of aberrations critical for MM prognostics and provides a feasible route for providing comprehensive genetics. Implementing lrWGS could provide more accurate clinical prognostics, facilitate genomic medicine initiatives, and greatly improve the stratification of patients included in clinical trials.

**KEY POINTS:**

- Linked-read WGS can be performed without DNA purification and allows for resolving the diverse structural variants found in multiple myeloma.
- Linked-read WGS can, as a stand-alone assay, provide comprehensive genetics in myeloma and other diseases with complex genomes.

## INTRODUCTION

Multiple myeloma (MM) is a hematological malignancy affecting terminally differentiated B lineage cells and is characterized by the accumulation of clonal plasma cells in the bone marrow ^1^. The disease has a complex genetic landscape thought to cause the clinical heterogeneity of the disease both in terms of treatment response and overall outcome ^2^. The introduction of novel treatments has significantly improved survival ^3^. However, despite these advances, the disease essentially remains incurable and patients with high-risk aberrations continue to display a poor outcome ^3–5^. Thus, identifying high-risk patients in the context of developing therapy regimens and understanding underlying disease biology to find novel venues for treatment remains critical to improve overall outcome.

Based on the primary genetic events, MM is largely divided into hyperdiploid (HRD) and non-HRD cases ^2,6^. In this division, HRD MM is characterized by multiple trisomies of odd-number chromosomes while non-HRD MM is associated with immunoglobulin heavy chain (IGH) translocations. The most common IGH translocations include t(4;14), t(11;14), t(6;14), t(14;16), and t(14;20) that, via colocalization of the strong IGH enhancers, cause the dysregulation of MMSET/FGFR3, CCND1, CCND3, MAF, and MAFB respectively. Traditionally, these primary aberrations have been investigated, together with common secondary events linked to poor outcome (including deletions of 17p, amplification of 1q21, and *MYC* translocations), in clinical routine using fluorescence in situ hybridization (FISH). Despite this seemingly simple dichotomy of initiating events, next-generation sequencing has revealed a complex landscape of genetic aberrations ^5,7-12^. In addition to the common initiating events, MM typically presents with a complex array of secondary genetic aberrations with numerous structural variants (SVs), copy number variations (CNVs), and single nucleotide variants (SNVs) affecting plasma cell differentiation, cell-cycle regulation, DNA repair and multiple signaling pathways ^2^. Major efforts are being made to exploit targetable aberrations, which together with high-throughput sequencing-based genomics open for the possibility of employing personalized medicine strategies for the treatment of MM ^2,13-15^.

Linked-read whole genome sequencing (lrWGS) is a developing technology that allows for the creation of synthetic long-reads from conventional short-read sequencing. Linked-read data is achieved by generating groups of reads or “read-clouds” originating from a single high-molecular-weight (HMW) DNA molecule, that all carry a common barcode linking them together ^16–19^. Mapping the read-clouds together and subsequently leveraging the existence of SNVs within the read-clouds allows for improved mapping and haplotype reconstruction of large phase blocks ^16^. The lrWGS data further provides the ability to identify and reconstruct SVs by analyzing read-clouds spanning chromosomal breakpoints. Several studies have effectively utilized lrWGS to identify and describe often complex SVs in various cancers and other diseases ^20–26^. This makes lrWGS an attractive option for providing comprehensive genetics in MM and other diseases with complex genomes.

Here we show that high-quality lrWGS can be generated directly on denatured FACS sorted cells without the need for prior DNA purification. Applying this to MM, we found that lrWGS reliably detected translocations and CNVs investigated by FISH. In addition, we show that lrWGS allows for identifying and resolving diverse structural events including private SVs with implications for the prognosis of the individual patient.

## MATERIALS AND METHODS

### MM patient samples and FACS sorting

Bone marrow samples were collected as part of the clinical routine from MM patients at Karolinska university hospital, Stockholm, Sweden. The collection and use of patient material was approved by the Swedish Ethical Review Authority (2014-526-31/3, 2019-02638 and 2020-00175) and performed in accordance with the Helsinki declaration. To analyze pure subsets, cells were FACS sorted using a FACS ARIAIIu (BD Biosciences) Propidium iodide (PI) was used to exclude dead cells. Using CD319 to compensate for the loss of CD138 on cryopreserved material ^27,28^, MM cells were FACS sorted as PI^-^CD14/CD16/CD11b^-^CD3^-^ CD319^+^CD38^+^. In addition, CD138, CD19, CD45, and CD56 were used to define the MM cells depending on expression. Myeloid and T-cells were FACS sorted as PI^-^CD14/CD16/CD11b^+^CD3^-^CD19^-^CD56^-^CD319^-^CD45^+^CD38^-^ and PI^-^CD14/CD16/CD11b^-^ CD3^+^CD19^-^CD56^-^CD319^-^CD45^+^CD38^-^ respectively.

### FISH

FISH was performed as previously described ^29^ using FISH probes targeting t(4;14), t(14;16), t(11;14), 1q21, 4p16, 6q21, 8p12, 9p21, 11q13, 13q14, 13q34, 14q32, 15q22, 16q23, 17p13 and 19q13.

### Linked-read WGS

To generate libraries, 200-240 cells were FACS sorted into 8μl PBS with 2% FCS, tubes were briefly pulse-spun, frozen on dry ice, and stored in −80°C until use. To denature the cells and make the genomic DNA accessible, 2μl 0.25N NaOH was added using narrow-bore tips and the sample was mixed by carefully flicking the tubes. After 5 min incubation at room temperature, 97.5μl of sample master mix (89.5μl Genome Reagent Mix, 3μl Additive A, 5μl Genome Enzyme Mix; Chromium Genome Reagent Kit v2 chemistry) was added to the denatured cells with a narrow-bore tip. Subsequently, the sample was mixed by gentle pipetting using a wide-bore tip. 90μl of the sample mix (containing ≈1.2ng DNA) was loaded onto the Chromium Genome Chip and the remaining steps were performed according to the manufacturer’s instructions (Manual CG00043 Rev B). Barcoded libraries were quantified using the Qubit dsDNA HS Kit (Invitrogen), pooled, and paired-end sequenced (2×150 cycles) using the Illumina platform (Novaseq, Illumina, San Diego, CA).

Data was processed using Long Ranger ^16^ and output files were used for downstream analysis. In brief: chromosome numbers were calculated using BarCrawler^30^ in combination with inhouse scripts; for selected samples, chromosome numbers were assessed also using CNVkit (v0.9.8)^31^, ASCAT (v2.5.2)^32^ and the Battenberg approach (v2.2.9)^33^; CNVs were called using FindSV^34^ and Long Ranger; Somatic variant calling was done using the nf-core/Sarek pipeline (v2.6)^35^ with Strelka2 (v2.9.10)^36^ as variant caller and a combination of VEP (v99.2)^37^ and SnpEff (v4.3t)^38^ for variant annotation; SVs were identified using GROC-SVs (v0.2.5)^39^ in combination with Long Ranger and called SVs were visually confirmed in Loupe and IGV^40^. For details see Supplemental Methods.

### RNAseq

Strand-specific RNAseq data was generated from FACS sorted MM cells as previously described ^41^ (for details see supplemental methods). Reads were mapped to hg38 using STAR (v 2.5.2b)^42^, reads in exons quantified using HOMER ^43^ and TPMs calculated using R.

### ChlPseq

ChlPseq was performed on 20k FACS sorted MM cells as previously described ^44^ using H3K27Ac antibodies (Diagenode Cat #C15410196, lot A1723-0041D/2) but with minor modifications on the preparation of input controls. Fastq files were mapped to hg38 using bowtie2 {Langmead2012} and downstream analysis performed using HOMER ^43^, DEseq2 ^45^ and R. For details see Supplemental Methods.

### Data sharing statement

Omics data is available through the SciLifeLab data repository (DOI: 10.17044/scilifelab.17049059). In compliance with Swedish law, sequencing data that could allow the identification of an individual is considered sensitive private information and therefore, data is under restricted access. Request for access can be made to the corresponding author.

## RESULTS

To investigate if lrWGS could be used to identify recurrent and private genetic aberrations in MM, we subjected a cohort of 37 MM samples to lrWGS (Table S1). All bone marrow samples were taken for diagnostic purposes (35/37 at diagnosis) and analyzed by an extended FISH panel. To ensure that analysis was performed on pure populations, all cells utilized were FACS sorted (Fig. 1A). In addition, germline controls were generated in 32/37 cases by performing lrWGS on primarily T-cells (Table S1). Prepared lrWGS libraries were sequenced to an average depth of 31.2x and 31.6x for MM and germline controls, respectively (Table S1).

**Figure 1.**
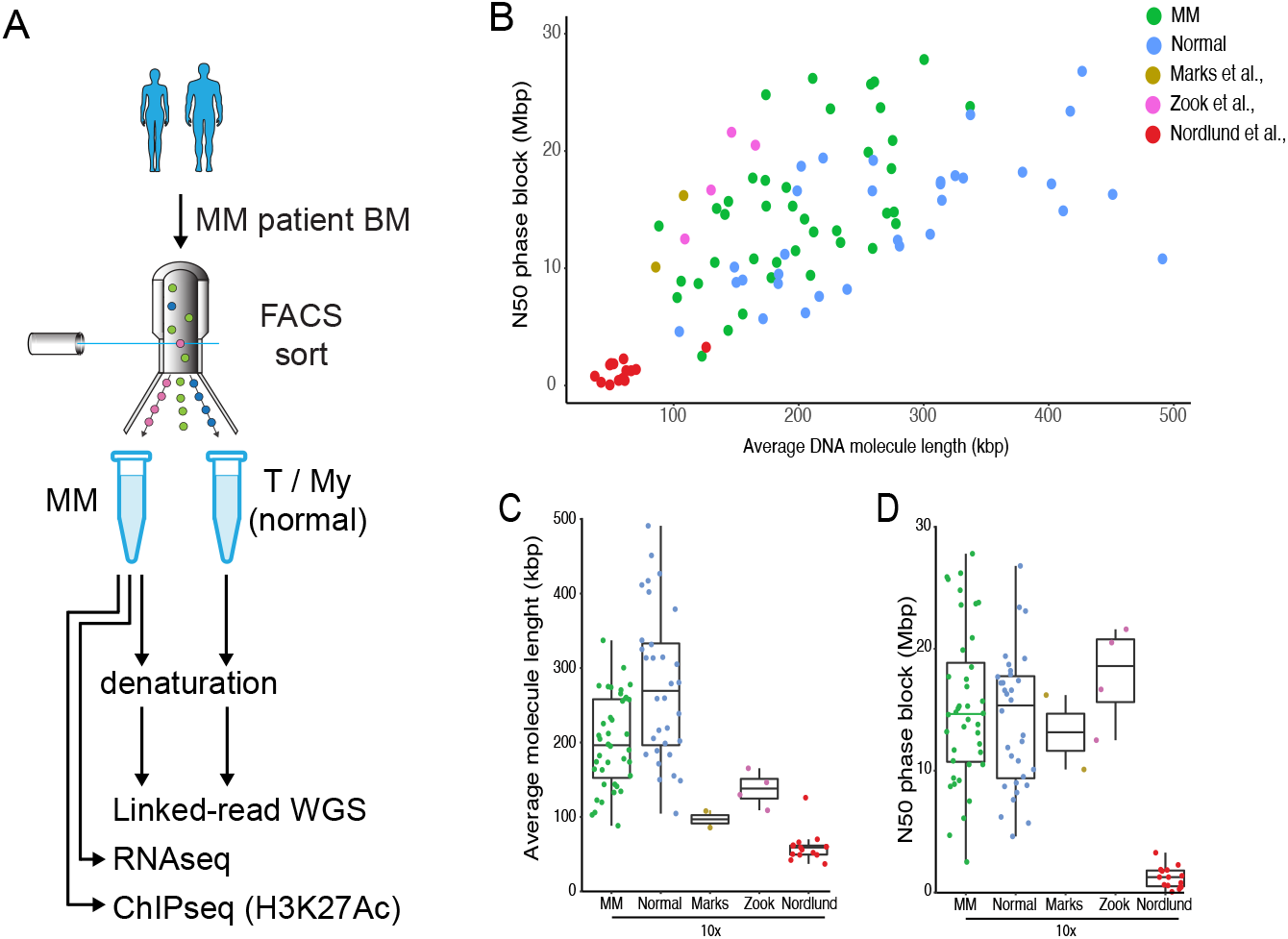
Linked-read whole-genome sequencing can be performed directly on denatured cells FACS sorted from diagnostic bone marrow samples. (A) Schematic overview of experimental setup. To perform genetic, transcriptional, and epigenetic characterization, multiple myeloma (MM) and normal cells (T, T-cells or My, myeloid cells) were FACS sorted from patient bone marrow (BM) samples. Linked-read WGS (lrWGS) libraries were prepared directly on denatured cells without prior DNA purification. Scatter plot (B) and box plots (CD) showing the N50 phase block size and average DNA molecule length of lrWGS libraries prepared using the indicated cell type and method. Quality control data from published lrWGS generated from prepared DNA (high-molecular weight or column purified) on the 10X platform are shown for comparison ^16,26,46^.

### High-quality lrWGS libraries can be prepared from low numbers of denatured cells

To fully utilize the lrWGS technology, libraries need to be generated from purified HMW DNA ^16,46^. To circumvent the need for HMW preparation, we established a protocol for preparing lrWGS libraries directly on cells denatured with sodium hydroxide. This allowed us to prepare lrWGS libraries on 200-240 FACS sorted cells (containing ≈1.2ng of DNA) on the Chromium Genome platform ^16^. We found that this simple protocol allowed for maintaining very large DNA molecules (median DNA fragment length of 216kbp), which was directly reflected in the size of the reconstructed phase blocks (median N50 of 14.8Mbp and median longest phase block of 60.2Mbp) (Fig. 1B-D, Table S1). The longest assembled phase block was 120.9 Mbp and constituted almost the entire q-arm of chromosome 4 (120.9Mbp/138.4Mbp). Thus, this phase block nearly reached the theoretical maximum length achievable without crossing centromeric regions. As expected, MM samples typically displayed poor phasing in areas with loss of heterozygosity (Fig. S1) but overall, the N50 phase block size of the MM samples was comparable to that of the normal controls (Fig. 1B-C). Comparing our lrWGS data to high-quality data sets generated from purified HMW DNA on the 10X platform ^16,46^, we found that our libraries were generated from significantly longer DNA molecules and maintained similar phase block sizes (Fig. 1B-D).

Overall, we conclude that performing lrWGS library preparation directly on denatured cells maintains DNA integrity and allows for generating high-quality lrWGS from very limited numbers of FACS sorted cells.

### Copy number variations identified by lrWGS and FISH are highly concordant

CNVs are a common feature of MM defining the HRD subgroup and high-risk patients ^3^. To identify CNVs, we calculated copy-numbers and computationally called CNVs. Comparing the chromosomal setup calculated from the lrWGS data to FISH, we found that chromosome numbers were correctly predicted in 35/37 (95%) MMs. In the two MM where prediction failed, no employed CNV caller (including FindSV^34^, CNVkit ^31^, ASCAT ^32^ and the Battenberg approach ^33^) accurately predicted their chromosome setup (data not shown). Nevertheless, correcting the copy-numbers based on chr4 FISH results allowed for visually identifying all (12/12) CNVs found by FISH (Fig. 2A). In the 35 patients where the chromosomal setup was accurately predicted from the lrWGS, 96% (143/149) of all CNVs identified by FISH – including all 1q amplifications - were concordantly called in the lrWGS data (Fig. 2A-B). These results are similar to those in prior studies comparing FISH with conventional WGS for identification of CNVs ^47,48^.

**Figure 2.**
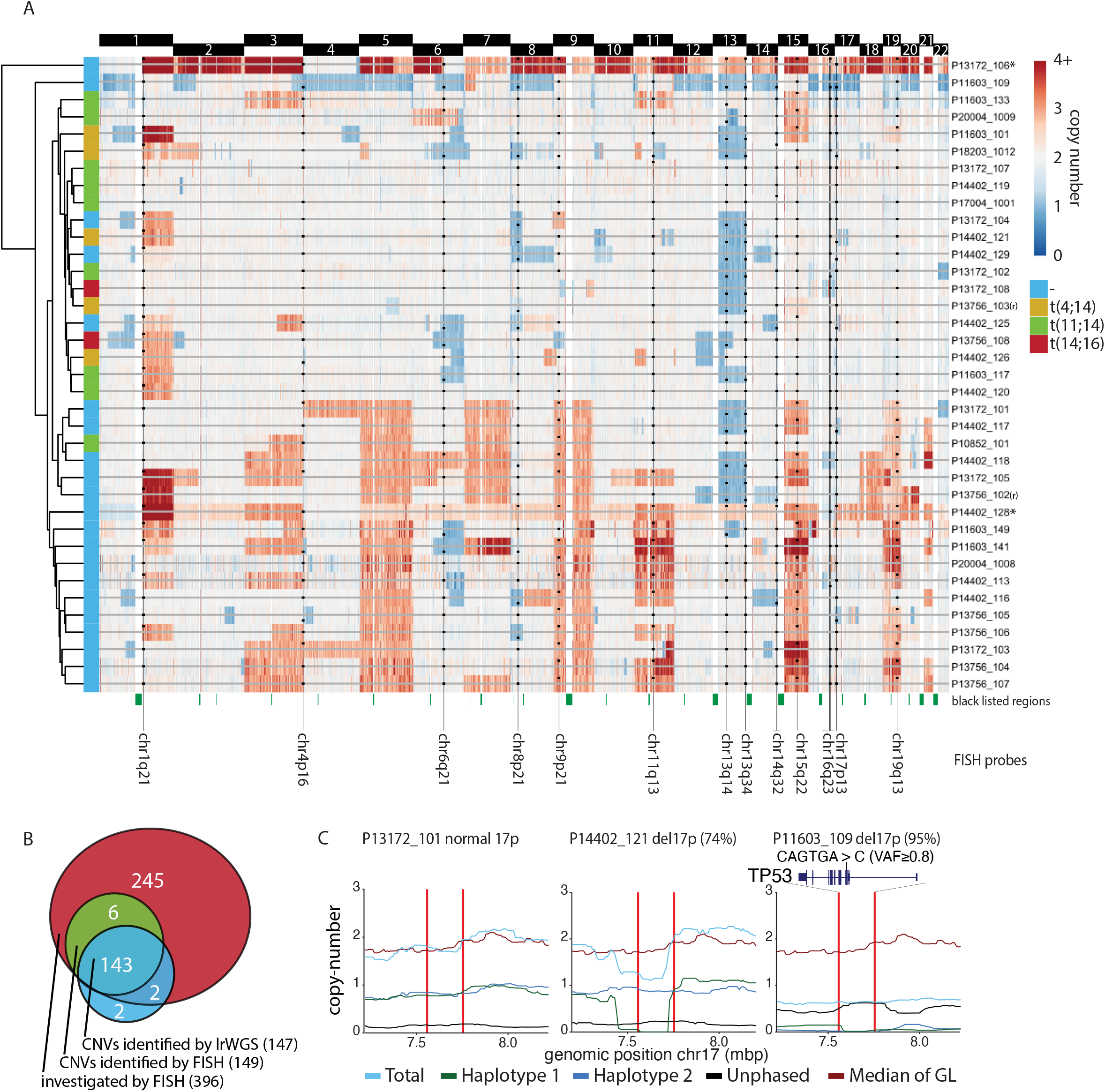
lrWGS allows for reliable detection of CNVs and identification of *“double hit” MM*. (A) Heatmap showing the copy-number across the somatic chromosomes (chr1-22) as calculated from lrWGS coverage. Copy-number was in two cases (indicated by *) corrected to the chr4 copy-number. Vertical black lines indicate genomic localization of FISH probes. Regions investigated by FISH are indicated by black dots with the position of the dot above or below the grey line indicating copy number gains or losses respectively. (B) Venn diagram showing the overlap between all regions investigated by FISH, CNVs identified by FISH and CNVs identified by lrWGS. (C) Copy-number of total, haplotype specific, and unphased reads in the area surrounding the TP53 locus. Median copy-number in the normal control samples (GL, germline) is shown for comparison. Vertical lines indicate the boundaries of the FISH probes used to identify chr17p CNVs. For P11603_109, the position of the acquired TP53 mutation is indicated.

Overall, this shows that lrWGS can reliably identify CNVs. However, our data argues that caution should be taken to verify chromosome numbers predicted by WGS in tumors with potentially widespread and complex subclonal copy-number changes.

### Identifying double-hit TP53 inactivation

Double-hit TP53 inactivation in MM is associated with poor outcome even with novel therapy ^3-5^. This prompted us to carefully investigate the existence of such events in our patient cohort. Haplotype specific chr17p deletions could readily be observed in the phased lrWGS data (Fig. 2D). Based on the computational calling of CNVs, we identified 7/8 (87.5%) patients with CNVs involving the *TP53* locus (Fig. S2). The 17p deletion with the lowest clonal involvement identified in the lrWGS data was found in 24% of cells by FISH (Fig. S2). In comparison, the undetected 17p deletion found by FISH was in only 14% of the cells. We next identified acquired mutations using Sarek ^35^ and analyzed sequence changes in genes recurrently mutated in MM ^9,11^ (Fig. S3A). We identified two patients with frame-shift mutations deleterious to TP53 function (Fig. S3A-B). The first was subclonal and appeared in a patient with an otherwise normal TP53 locus (P13756_103). In contrast, the second mutation was near clonal (present in ≥80% of reads in patient P11603_109) and co-occurred with a clonal 17p deletion (Fig. 2D, right) suggesting that this patient presented with high-risk double-hit TP53 MM at diagnosis.

### The primary MM IGH locus translocations are readily identifiable by lrWGS

The analysis of read-clouds spanning regions on different chromosomes allows for accurate identification and reconstruction of SVs (for examples and an in-depth explanation of data interpretation see Fig. S4). We utilized Long Ranger ^16^ and GROCS-SVs ^39^ to call inter-chromosomal SVs. This approach identified 198 SVs across all samples (Fig. 3A and Table S2). No significant differences in the number of SVs per sample were found between HRD and non-HRD cases though three cases of HRD MM had no detectable SVs (Fig. 3B).

**Figure 3.**
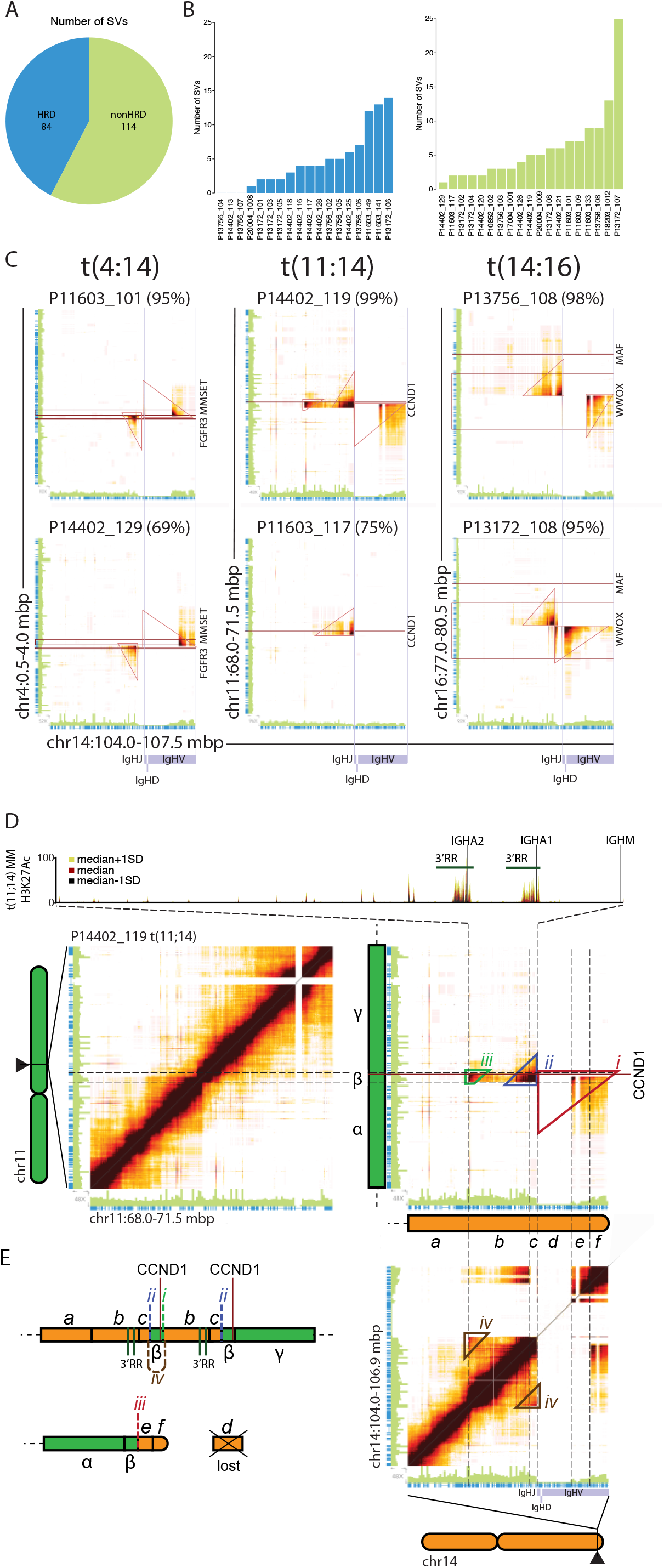
Breakpoint spanning read-clouds allow for identification and reconstruction of IGH translocation events. (A) Total number of SVs involving different chromosomes identified in HRD and non HRD MM. (B) Number of SVs involving different chromosomes in HRD (left) and non HRD MM (right). (C) Heatmaps displaying the number of read-clouds shared between the IGH locus (chr14) and breakpoint regions on chr4 (MMSET), chr11 (CCND1) and chr16 (MAF). Barcode overlap between indicated regions were drawn by Loupe with the yellow-red-black color scale indicating progressively higher barcode overlaps. Sequencing coverage (green bars) and the positions of coding regions (blue bars) is indicated on the axis of the heatmap. The percentage of cells found by FISH for each translocation is indicated in parenthesis above each heatmap. Horizontal (red) and vertical (purple grey) lines mark the position of indicated genes and IGH VDJ regions respectively. (D) Read-cloud overlap on chr11 (top left), chr11 to chr14 (top right) and chr14 (bottom right) in P14402_119. Track shows the H3K27Ac signal (median±SD) in t(11;14) MM. Features (*i-iv*) used to resolve the structure of the derivate locus are indicated by colored triangles. (E) Schematic representation of the derivate chromosomes in P14402_119 as determined by lrWGS.

To identify IGH translocations, we visualized the common t(4;14), t(11;14), and t(14;16) breakpoint regions using Loupe. We found that the translocations could readily be identified visually through the enrichment of breakpoint spanning read-clouds (Fig. 3C and S5). The enrichment was highly specific to the individual IGH translocation events and clearly distinguishable from background signals (Fig. S5). Among the computationally called SVs (Table S2), we found 16 MMs with IGH translocations involving the *MMSET, CCND1* or *MAF* locus. Overall, 16/17 (94%) of the IGH translocations identified by FISH were found by lrWGS and no false-positive calls were made. The unidentified t(11;14) was subclonal (28% of cells) and read-clouds supporting the existence of the SV were identified upon visual inspection (Fig. S5).

Together, this suggests that lrWGS allows for accurately identifying SVs including the common IGH translocations.

### t(11;14) are associated with amplification of the 3’RR enhancer regions

In line with the notion that the IGH translocation events often occur as a result of erroneous class-switch recombination ^2^, the IGH translocations were localized on the centromeric side of the IGHJ-regions (Fig. 3C and S5). Looking specifically at the t(4;14) and t(14;16) translocations we found that they were all reciprocal with the chr14 breakpoint occurring between the Eμ and the 3’RR enhancer regions (flanking the IGHA1/2 region) (Fig. S6 and S7) ^49^. In contrast, almost half (4/9) of the t(11;14) MMs presented with patterns of read-cloud clusters indicative of more complex events (Fig. 3C and S5). The t(11;14) cases with simple reciprocal or non-reciprocal translocations (5/9 cases), resulted in the 3’RR regions being juxtaposed with the *CCND1* locus (Fig. S8). Looking specifically at the t(11;14) cases with multiple read-cloud clusters, three of the cases (P14402_119, P17004_1001, and P13172_102) shared a similar pattern with three overlapping clusters (Fig. S5) consistent with additional events occurring in conjunction with a reciprocal translocation. Analyzing the read-cloud overlaps of the clonal t(11;14) in P14402_119, we found that the pattern was consistent with a reciprocal translocation (Fig. *3Di-ii*) co-occurring with a focal amplification of the translocation breakpoint region on the CCND1 carrying derivative chromosome (Fig. *3Dii-iv* and E). The results were supported by the copy-number gains on chr11 (Fig. 3D, β region) and chr14 (Fig. 3D, b-c region) as well as the shared read-clouds between centromeric (5’) and telomeric (3’) ends of the amplified region on chr14 (Fig. *3Div*, b-c region) and the high read-cloud overlap in the middle event (Fig. 3D*ii*). The two other cases with the same pattern of read-clouds involving overlapping regions (P17004_1001 and P13172_102) could be resolved in the same manner. Looking at the fourth case (P14402_120) - which had a different pattern with two (in terms of genomic location) partially overlapping read-cloud clusters - we concluded that this pattern represented a reciprocal translocation with duplicated flanking regions on either side of the translocation breakpoint (Fig. S9). Overlapping the involved regions with H3K27Ac (marking active gene regulatory elements) ChIPseq data, we found that the 3’RR regions were within the involved regions in all four cases (Fig. 3D and S10A). As expected, this was associated with a very high expression of *CCND1* (>1000 TPM in all four patients) (Fig. S10B).

Taken together, this suggests that t(11;14) events are frequently associated with either focal or breakpoint flanking amplifications affecting the *CCND1* locus and the 3’RR of the IGH locus.

### lrWGS allows for detecting and resolving diverse SVs affecting the MYC locus

*MYC* translocations have been shown to be associated with poor outcome ^10,50^. SVs involving the *MYC* locus are often complex involving focal amplifications, insertions, and translocations that are thought to dysregulate *MYC* expression by positioning the gene in proximity with strong enhancer elements^10,50,51^. Relying on the computational identification of SVs (Table S2), we found SVs affecting the *MYC* locus in 9/37 (24%) patients with the IGL region being the only recurrently involved locus (2/9 cases). The identified SVs were supported by distinct read-cloud clusters (Fig. 4A) that could easily be distinguished from the background (Fig. S11). The majority of the identified *MYC* SVs were simple translocations (5/9) (Fig. 4Ai). Looking closer at the remaining cases with overlapping read-cloud clusters, we concluded that these cases represented focal amplifications with templated insertions (Fig. 4Aiii and B) or reciprocal translocations with duplications of the breakpoint flanking regions (Fig. 4Aii and C). Both kinds of SVs have previously been described for the *MYC* locus^50,51^. As expected, identified SVs involved either the region containing the *MYC* and *PVT1* promoters (6/9 cases) or the area just telomeric of the *PVT1* gene (3/9 cases) known to harbor elements influencing the *MYC* gene^52^. All three SVs affecting the *PVT1* flanking enhancer region resulted in the juxtaposition of this element with myeloma-active gene regulatory elements from another chromosome through insertion (P14402_118 and P11603_101) (Fig. 4B and S12A) or by reciprocal translocation (P13756_108) (Fig. 4C). Similarly, the translocations affecting the *MYC*/*PVT1* promoter area in most cases resulted in the *MYC* locus being juxtaposed with myeloma-active elements from various genomic locations including the JCHAIN and IGL loci (Fig. S12A-B). Overall, MM with *MYC* SVs expressed higher levels of *MYC* than MM cases with a normal *MYC* locus but the difference was not statistically significant (Fig. S12C)^10^.

**Figure 4.**
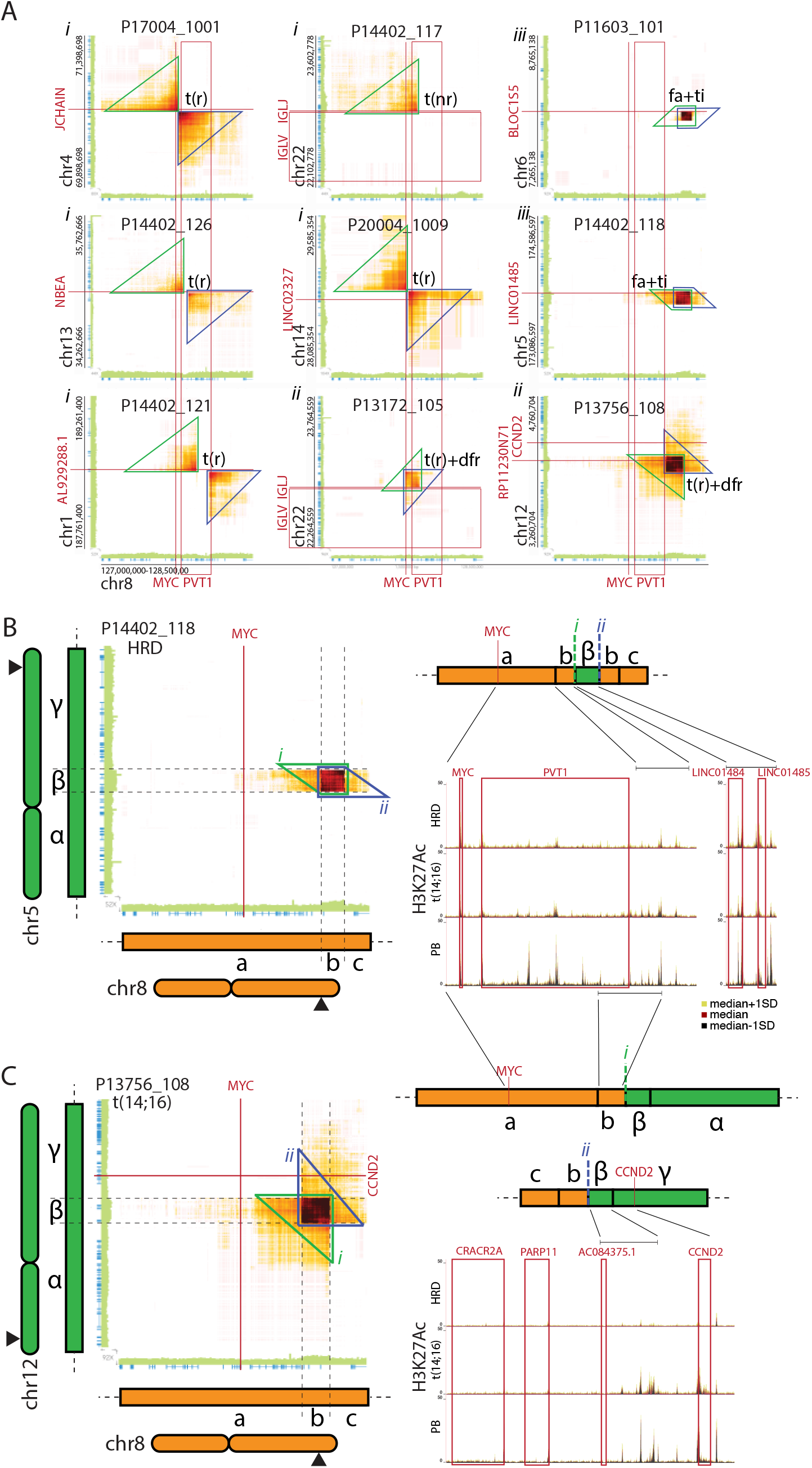
Diverse structural variants affecting the MYC locus can be identified by lrWGS. (A) Heatmaps displaying the number of read-clouds shared between the MYC locus and indicated genomic locations. Barcode overlap between indicated regions were drawn by Loupe with the yellow-red-black color scale indicating progressively higher barcode overlaps. Sequencing coverage (green bars) and the positions of coding regions (blue bars) is indicated on the axis of the heatmap. Data from all patients (n=9) with detected SVs affecting the genomic MYC regions are shown. The type of SV is indicated next to the read-cloud clusters supporting the existence of the SV: t(r), reciprocal translocation; t(nr), non-reciprocal translocation; t(r)+fa, reciprocal translocation with focal amplification; and t(r)+dfr, duplication of breakpoint flanking regions. (B-C) Read-cloud overlap between the MYC locus on chr8 and chr5 in P14402_118 (B) or chr12 in P13756_108 (C). Features (*i-ii*) used to resolve the structure of the derivate locus are indicated by colored triangles. Schematic representations of the derivate chromosome regions are shown on the right. Tracks show the H3K27Ac signals (median±SD) for the involved regions in the indicated MM subtypes and plasma blasts (PBs).

Together, this shows that the *MYC* locus is affected by a diverse spectrum of SVs in MM and that these can be detected and structurally resolved using lrWGS.

### Private structural variants cause the overexpression of MAFA and MAP3K14

To determine if we could find additional SVs that may represent potential primary or high-risk variants, we investigated whether we could identify private SVs affecting loci previously found to be translocation partners of the IGH locus ^2,50,53^. This identified a t(1;8) in P13172_104 involving the *MAFA* locus and a t(6;17) in P13172_101 involving the *MAP3K14* locus.

Patient P13172_104, had two SVs involving chromosomes 1 and 8 with the second juxtaposing the *MAFA* and TENT5C loci (Fig. 5A-C). Analysis of the read-clouds in the breakpoint regions suggests that the SV involving *MAF* was clonal (Fig. S13A-B). Looking closer at the breakpoints, we found that the *MAFA* gene was juxtaposed with a strong MM-active enhancer region normally flank the *TENT5C* gene (Fig. 5C). H3K27Ac ChIPseq confirmed that both the *TENT5C* flanking enhancer and *MAFA* proximal elements were highly active in MM cells from P13172_104. Hence, this strongly suggests that the TENT5C enhancer causes activation of the MAFA gene. In line with this, we found very high *MAFA* expression (approximately 1000 TPM) specifically in the MM cells from P13172_104 (Fig. 5D). Given that MAFA is part of the Maf transcription factor family, we next wanted to assess if the *MAFA* overexpression resulted in a gene-regulatory landscape similar to that in t(14;16) *(MAF* overexpressing) MM. Looking at MM subgroup-specific H3K27Ac signals, hierarchical clustering of all the samples revealed that P13172_104 clustered with the t(14;16) MM and shared H3K27Ac features with the *MAF* overexpressing group (Fig. 5E). Hence, the private t(1;8) translocation in P13172_104 results in the overexpression of MAFA and the establishment of a MAF-type gene regulatory landscape.

**Figure 5.**
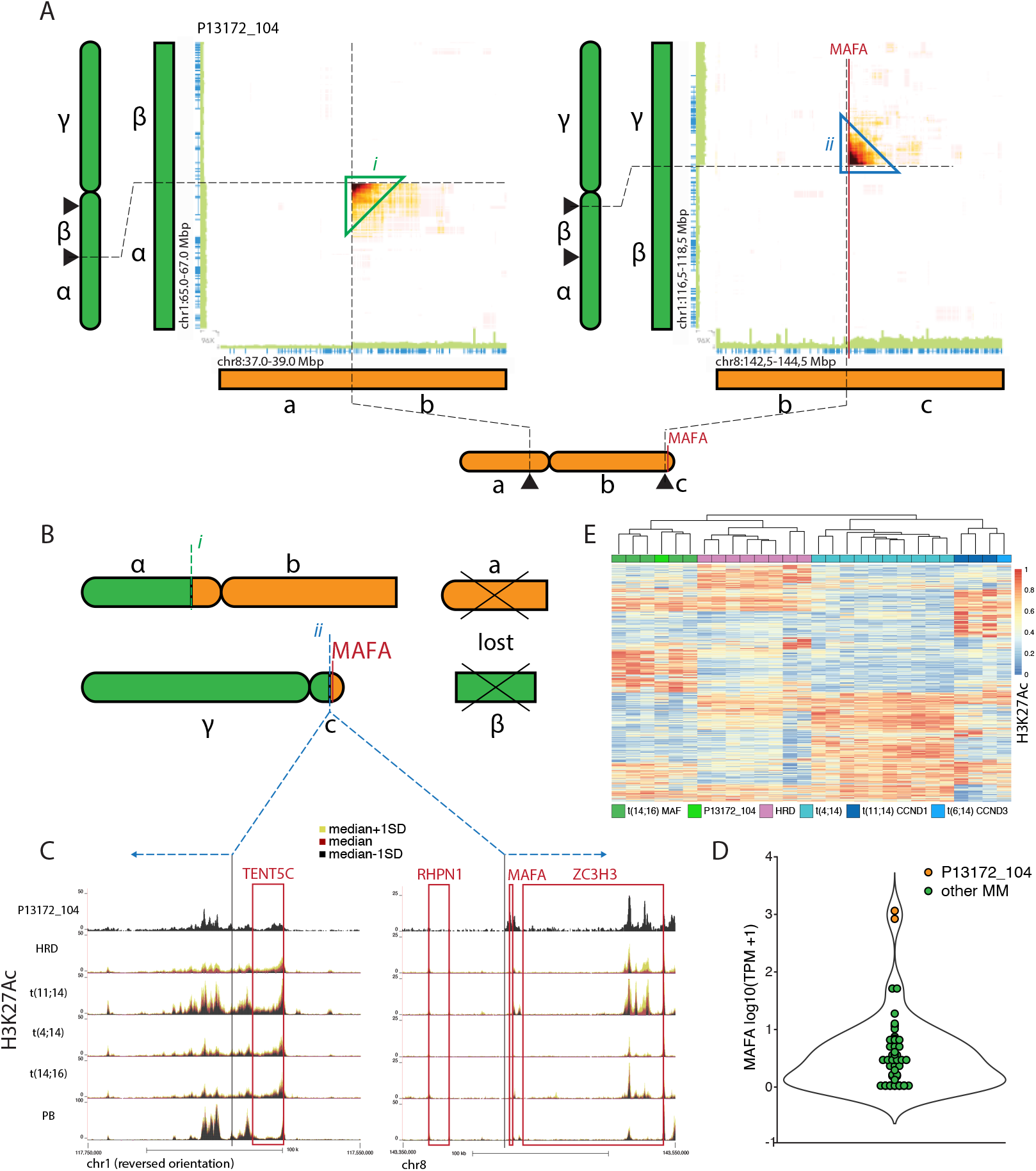
A private SV cause massive MAFA overexpression and reprogramming to MAF-type MM. (A) Heatmaps displaying the number of read-clouds shared between indicated regions on chr1 and chr8. Barcode overlap between indicated regions were drawn by Loupe with the yellow-red-black color scale indicating progressively higher barcode overlaps. Sequencing coverage (green bars) and the positions of coding regions (blue bars) is indicated on the axis of the heatmap. (B) Schematic representation of the derivate chromosomes. (C) Tracks showing the H3K27Ac signal (median±SD) in the regions involved in the t(1;8). (D) Expression of MAFA. (E) Row normalized hierarchically clustered heatmap of H3K27Ac marked regions showing differential signals between MM subtypes.

The SV affecting the *MAP3K14* locus structurally constitutes a focal amplification on chr17 carrying a templated insertion of a small chr6 region (Fig. 6A-B). Analysis of read-cloud overlaps indicates that the SV was subclonal (Fig. S13C). Analyzing the involved regions, we found that the SV juxtaposed an enhancer region normally flanking the BMP6 gene with the amplified region on chromosome 17 (Fig. 6B). The latter contained part of the *MAP3K14* gene but lacked the 3’ most exon, indicating that the amplification does not result in two functional copies of the gene. In line with the *BMP6* flanking element dysregulating the MAP3K14 gene, expression was high (>300 TPM) in P13172_101 and almost 10-fold higher than any other investigated MM (Fig. 6C). With *MAP3K14* functioning as an activator of *NF-KB,* this could suggest that the MM subclone carrying this SV could present with elevated or constitutive NF-KB signaling ^54^.

**Figure 6.**
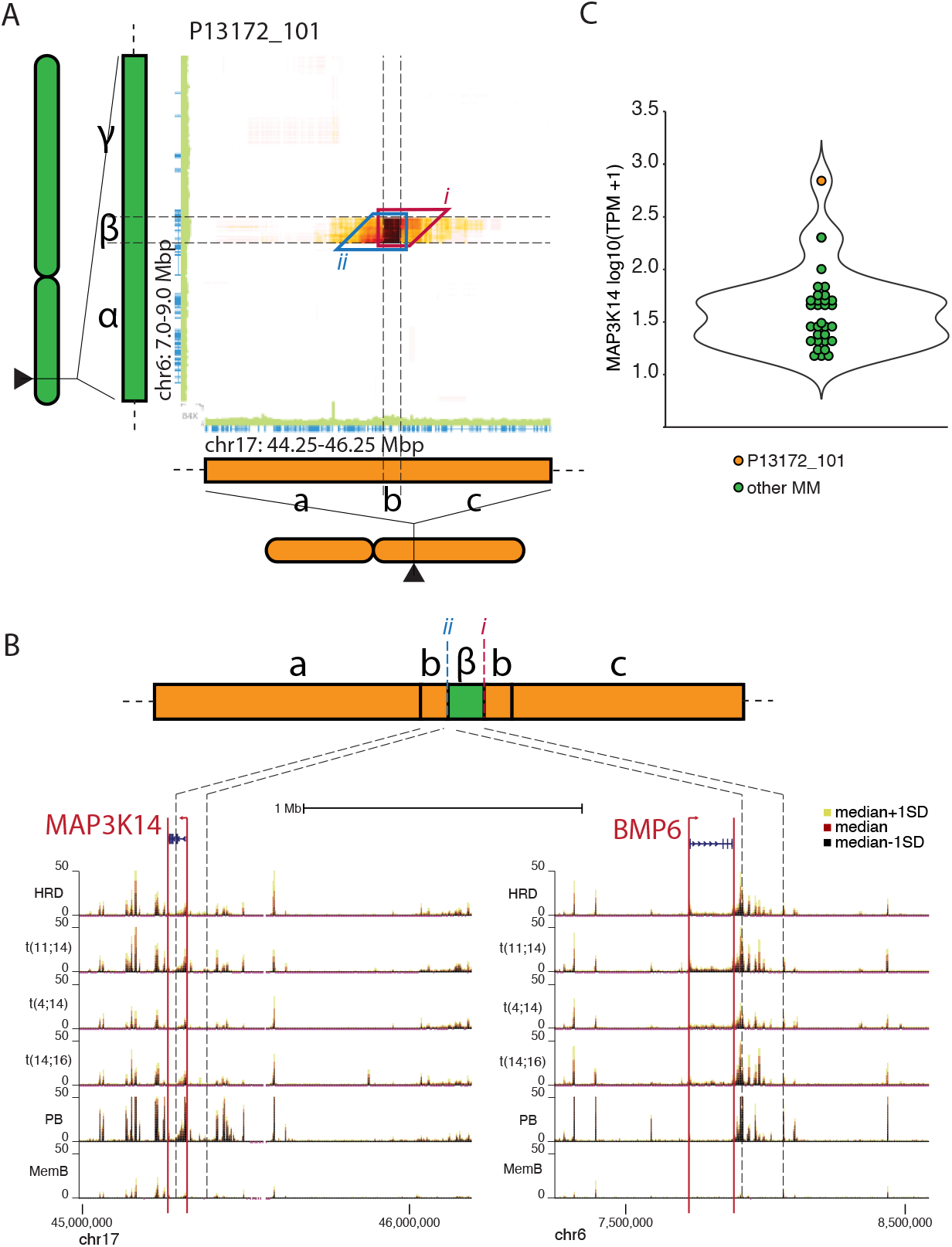
The insertion of the BMP6 flanking enhancer region causes the overexpression of MAP3K14. (A) Heatmaps displaying the number of read-clouds shared between indicated regions on chr6 and chr17. Barcode overlap between indicated regions were drawn by Loupe with the yellow-red-black color scale indicating progressively higher barcode overlaps. Sequencing coverage (green bars) and the positions of coding regions (blue bars) is indicated on the axis of the heatmap. (B) Schematic representation of the derivate chromosome and tracks showing the H3K27Ac signal (median±SD) in the regions involved in the SV. (C) Expression of MAP3K14.

## DISCUSSION

MM remains an incurable disease with poor outcome for most patients. Therefore, developing strategies to provide comprehensive genetic characterization in clinical routine is of paramount significance to understand the disease at the level of individual patients and for establishing personalized medicine programs aiming to improve outcome and survival.

Here we performed a proof-of-principle study on 37 MM patients and demonstrate that lrWGS can be used to provide comprehensive genetics in MM. Compellingly, the lrWGS provided accurate and straightforward identification of the recurrent IGH and IGL translocations through simply visualizing read-clouds spanning common breakpoint regions. Given the specificity of the read-cloud clusters, these SVs can be rapidly identified even without computational analysis (apart from base processing) or germline controls. In a clinical setting, this is attractive, as it would allow for making a quick screen of common SVs before performing more time-consuming holistic analysis. As exemplified by the SV affecting both the IGH and *MYC* loci, analysis of the read-cloud clusters can identify and resolve diverse SVs in a simpler and more accurate way than possible with FISH or conventional WGS. Overall, given the high concordance between the lrWGS and FISH, this suggests that lrWGS can function as a stand-alone assay that can alleviate the need to complement sequencing-based analysis with optical mapping^55,56^ or other technologies aimed at mapping SVs.

Taking advantage of the possibility to holistically analyze genetic aberrations in the lrWGS data, we identified both recurrent and private genetic events associated with a poor outcome that would have remained undetected in the current clinical routine genetics. Among the recurrent variants, we found both cases with t(8;22) MYC-IGL translocations and high-risk double-hit TP53 inactivation. Interestingly, we could also find private structural variants causing the overexpression of genes previously shown to be translocation partners of the IGH locus *(MAP3K14* and *MAFA*). In particular, the t(1;8) involving the MAFA gene is notable as this SV likely represents an alternative primary event that would be the molecular equivalent of t(14;16) deregulation of *MAF* with a similar patient risk profile and innate resistance to proteasome inhibitors^57^. Interestingly, we also found that t(11;14) translocations often carried duplications of the 3’RR and CCND1 regions. The fact that the t(11;14) MMs analyzed were randomly included suggests that this is a rather common phenomenon, which to our knowledge has not been previously described.

Nominally, lrWGS can be performed on very limited amounts of cells (≤1000 or 1-5ng of DNA) but the need for HMW DNA preparations has effectively barred this. Circumventing the need for DNA preparation – as done in this study – to fully take advantage of the low requirements on input material has interesting implications for how to perform routine genetics on MM and other hematological malignancies. By simply running diagnostic flow cytometry on FACS sorters lrWGS based genetics can be performed on any defined tumor or normal cell population. Given the low number of cells needed, this could even be performed on samples where tumor cells are expected to be rare or low in number, including samples from fine needle biopsies or samples taken for minimal residual disease analysis. This approach would simultaneously allow for performing RNAseq and other assays on the pure target cell population in parallel with genetics. The more recent introduction of more automated FACS sorters makes this kind of analytical workflow feasible to implement in most diagnostic flow cytometry laboratories.

While the removal of the Chromium Genome from the market undoubtedly has hampered the wider introduction of lrWGS, several other platforms have emerged to take its place, including single-tube long fragment read (stLFR)^58^, transposase enzyme-linked long-read sequencing (TELLseq)^59^ and droplet barcode sequencing (DBS) ^19^. While larger validation studies will need to be performed to demonstrate that these effectively can replace the 10X platform, the simple workflow of the TELLseq protocol makes this an interesting candidate for a potential clinical implementation.

In summary, lrWGS can with relative ease identify and resolve the structure of diverse SVs making it an attractive solution for providing comprehensive genetics in MM and other hematological malignancies with complex genomes. Furthermore, utilizing FACS sorting in diagnostics could facilitate lrWGS of pure tumor cells as well as molecular characterization on both the expression and epigenetic level. This could provide a powerful platform for understanding underlying genetics and disease mechanisms in individual patients.

## Supporting information

10xMM_Supplemental

Table_S2_Translocations

## AUTHOR CONTRIBUTIONS

RM proposed the studies and devised strategies for performing lrWGS without DNA preparation. LPP, NF, CGr, and RM planned the study. AW biobanked patient samples and performed FISH. CGr, CGu and FTL performed proof-of-principle lrWGS experiments. RAO and PE performed initial analysis on proof-of-principle experiments. NF, JH, CGr, AK and RM FACS sorted cells. CGr, CGu and FTL prepared 10X lrWGS libraries. LPP analyzed lrWGS data. JE, MK and AL assisted in establishing analysis pipelines. CGu performed ChIPseq. JH analyzed ChIPseq data. CGu and NF performed RNAseq. LPP and NF analyzed RNAseq data. PE, AL, and RM supervised the study. LPP and RM integrated data. LPP and RM wrote the manuscript with input from the other authors. All authors reviewed the manuscript before submission.

## ACKNOWLEDGEMENTS

We would like to thank: Hareth Nahi for providing patient material and basic patient information; the Centre for Cellular Analysis for assistance with cell sorting; the Swedish National Infrastructure for Computing (SNIC) at Uppmax for computational resources; National Bioinformatics Infrastructure Sweden (NBIS) for support and mentoring; the National Genomics Infrastructure for sequencing service; Aron Skaftason for input on variant calling; and Joakim Dillner for access to sequencing equipment. This work was supported by the Swedish Cancer Foundation, Swedish Research Council, King Gustav V Jubilee Fund, the Stockholm County Council, the Swedish Foundation for Strategic Research, and the Knut and Alice Wallenberg Foundation.

