## Supplementary material for "Linked-read whole-genome sequencing resolves common and private structural variants in multiple myeloma": 10xMM_Supplemental

#### SUPPLEMENTAL METHODS & FIGURES

#### SUPPLEMENTAL METHODS

##### FACS antibodies

The majority of samples were stained with an antibody panel containing: CD14 PE-Cy5 (61D3, eBiosciences), CD16 PE-Cy5 (3G8, BD), CD11b PE-Cy5 (ICRF44, BD), CD56 BV421 (HCD56, BioLegend), CD3 Alexa700 (UCHT1, BD), CD19 PeCy7 (HIB19, BioLegend), CD38 BV605 (HB7, BD), CD45 BV786 (HI30, BD), CD138 APC (MI50, BioLegend) and CD319 PE (162.1, BioLegend).

##### Generation of strand-specific RNAseq

500 or 1000 cells were FACS-sorted into 5 or 10μl Single cell lysis solution (SCLS) respectively (Single cell lysis kit Invitrogen, cat# 4458235) with DNAse I according to manufacturer's instructions with the exception that the lysis reaction was incubated for 15min at RT before adding stop solution. For a few samples, 20k cells were instead sorted into buffer RLT with β-mercaptoethanol and total RNA extracted using RNeasy Micro Kit (Qiagen, cat#74004) with on-column DNase I treatment according to manufacturer's instructions. Double stranded cDNA was prepared from 500 lysed cells in SCLS or the RNeasy purified RNA from 7.5-10k cells using NEBNext Ultra II RNA First Strand Synthesis (NEB cat#E7771) and NEBNext Ultra II Directional RNA Second Strand Synthesis (NEB Cat# E7550) modules according to manufacturer's instructions. In majority of samples, QIAseq FastSelect rRNA HMR kit (Qiagen cat# 334386) was utilized (according to manufacturer's instructions) to minimize rRNA contribution to the final library. Stranded libraries were subsequently prepared by adding custom made i5 containing Tn5 (transposase) and tagmentation buffer (50mM tris Acetate, 25mM Mg Acetate, 50% dimethylformamide) to double stranded cDNA sample. After 5 min at 55°C, the reaction was stopped by incubating with 5μl 1% SDS for 5min at room temperature and samples were purified using AmPure XP beads at a ratio sample:beads of 1:1.3 and eluted in water. Introduction of indexes through oligo replacement [1] was carried out by adding custom made i7 replacement oligos and gap fill buffer (165mM Tris acetate, 330mM potassium acetate, 50mM Mg acetate, 2.5mM DTT, 1mM beta-NAD and 1.25mM each of dATP, dCTP, dGTP and dTTP). After 30min incubation at 37°C, gap fill enzymes were added (1U Sulfolobulus DNA Polymerase IV NEB cat# M0327S and 10U E Coli DNA Ligase NEB cat# M0205L) and incubation continued for an additional 30min at 37°C. Samples were purified using AmPure XP beads at a ratio sample:beads of 1:1.25 and eluted in 12μl water. Libraries were subsequently PCR amplified using Phusion HF PCR Master Mix (Thermo Scientific) with the following PCR program: 95°C 2 min; followed by 16 cycles of 94°C 10 s, 60°C 30 s and 72°C 1 min. Library cleanup was done using AmPure XP beads at a ratio sample:beads of 1:0.9. DNA concentrations in purified samples were measured using the Qubit dsDNA HS Kit (Invitrogen). Barcoded libraries were pooled and paired-end sequenced (2×41 cycles) using the Illumina platform (NextSeq500, Illumina, San Diego, CA).

##### ChIPseq and analysis

ChIPseq was performed as previously described with minor modifications to the preparations of input controls [2] was added to each tube. Input control libraries were amplified together with ChIPseq libraries using the following PCR program: 72°C 5 min (adapter extension); 95°C 5 min (reverse crosslinking and denaturation); followed by 11 cycles of 98°C 10 s, 63°C 30 s and 72°C 3 min. After PCR amplification, library cleanup was done using Agencourt AmPureXP beads (Beckman Coulter) at a PCR mix to bead ratio of 1:1. Barcoded libraries were pooled and single-end sequenced (50 cycles) using the Illumina platform (NextSeq500, Illumina, San Diego, CA)

Fastq files were mapped to hg38 using bowtie2, identification of peaks and quantification of reads in peaks was done using HOMER [3]. Identification of elements displaying significantly different signals between different genetic MM subgroups (HRD, t(4;14), t(11;14) and t(14;16)) was done using DESeq2 [4]. Peaks with a fold change >2 and with an adjusted p-value <0.05 were considered significantly changed. The row-normalized values for the 250 most significant peaks in each comparison were plotted using R. A combination of newly generated and previously published H3K27Ac ChIPseq data [5] was used in the analysis. For visualization, median ($\pm$ standard deviation) H3K27Ac coverage for each MM genetic group was calculated and visualized in UCSC genome browser [6] using a multiWig hub.

**Analysis of linked-read WGS data**

*Identification of CNVs*

To calculate and visualize copy-number, Long Ranger [7] files were processed using BarCrawler[8] and the output subsequently processed by an in-house developed script that transforms read coverage into chromosome numbers. In brief, BarCrawler output was binned into 500kb windows and the median total, unphased and haplotype-specific coverage per chromosome was calculated. We considered a chromosome to have two copies when the median coverage per haplotype on the chromosome was ≥0.47x (median total coverage on the chromosome minus median unphased coverage on the chromosome) and the standard deviation of median coverage on the chromosome < 1.5x the median standard deviation of total coverage of all chromosomes. The median coverage per base on the chromosomes with two copies were divided by two to obtain a coefficient subsequently used to convert coverage to copy-number for all chromosomes. For selected samples, chromosomal copy-numbers were assessed using CNVkit (v0.9.8)[9], ASCAT (v2.5.2)[10] and the Battenberg approach (v2.2.9)[11]. CNVs were called using FindSV[12] and Long Ranger. CNVs >30kb overlapping with regions investigated by FISH were further confirmed by visualizing the haplotype specific coverage (10kb bins).

*Identification of somatic variants from lrWGS tumor/normal pairs*

Somatic variant calling was done using the nf-core/Sarek pipeline v2.6 [13]. In brief, the BAM files containing the Long Ranger mapping information were used as input for the Sarek pipeline, Strelka2 (v2.9.10)[14] was used for variant calling (tumor-normal subtraction analysis) and a combination of VEP (v99.2)[15] and SnpEff (v4.3t)[16] was used for variant annotation. Variants in genes recurrently mutated in MM [17,18] with a PASS filter (from Strelka2), having a moderate or high predicted impact on gene function according to either variant annotation tool and having a GnomAD allele frequency <1% were considered to effect gene function.

*Identification of SVs*

SVs involving different chromosomes were identified using GROC-SVs (only considering tumor lrWGS data) and Long Ranger. To exclude obvious artifacts, identified SVs were visually assessed using Loupe (10x Genomics, Loupe Browser 2.1.2). SVs called by Long Ranger and/or GROC-SVs that were not within blacklisted regions and passed visual inspection (using Loupe and IGV [19] to exclude artifacts) were considered valid.

Fig. S1

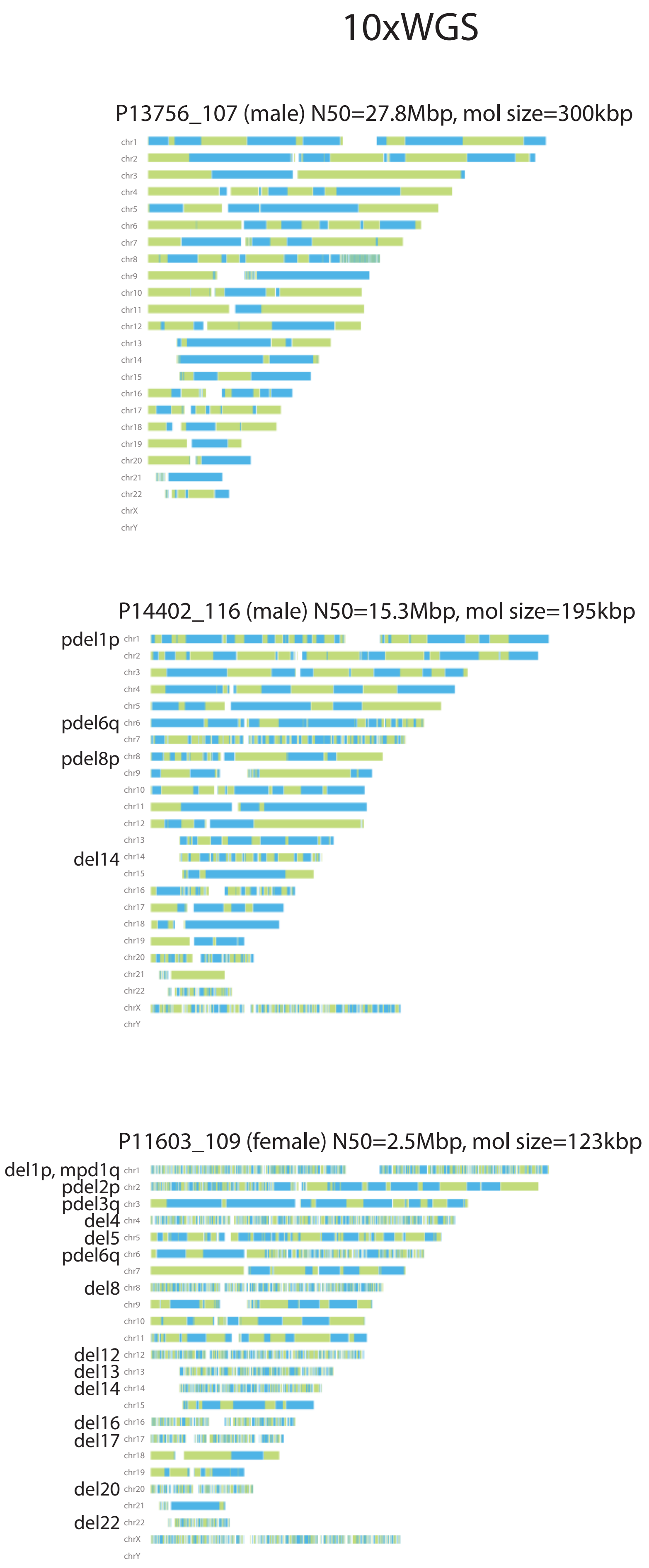

10xWGS
P13756_107 (male) N50=27.8Mbp, mol size=300kbp
chr1
chr2
chr3
chr4
chr5
chr6
chr7
chr8
chr9
chr10
chr11
chr12
chr13
chr14
chr15
chr16
chr17
chr18
chr19
chr20
chr21
chr22
chrX
chrY
P14402_116 (male) N50=15.3Mbp, mol size=195kbp
pdel1p
pdel6q
pdel8p
del14
chr1
chr2
chr3
chr4
chr5
chr6
chr7
chr8
chr9
chr10
chr11
chr12
chr13
chr14
chr15
chr16
chr17
chr18
chr19
chr20
chr21
chr22
chrX
chrY
P11603_109 (female) N50=2.5Mbp, mol size=123kbp
del1p, mpd1q
pdel2p
pdel3q
del4
del5
pdel6q
del8
del12
del13
del14
del16
del17
del20
del22
chr1
chr2
chr3
chr4
chr5
chr6
chr7
chr8
chr9
chr10
chr11
chr12
chr13
chr14
chr15
chr16
chr17
chr18
chr19
chr20
chr21
chr22
chrX
chrY

**Figure S1. Deletions are associated with poor phasing and shorter phase blocks.** Phase block structure of the samples with the highest (top), representative intermediate (middle) and shortest (bottom) N50 phase block size are shown. Phase blocks were drawn by Loupe in green and blue, where alternate colors illustrate phase block boundaries. Patient number, N50 and average molecule size are given for each sample above the graphical representations. Known copy number variations are indicated on the left of each chromosome (del, deletion; pdel, partial deletion; and mpd, monoparental disomy).

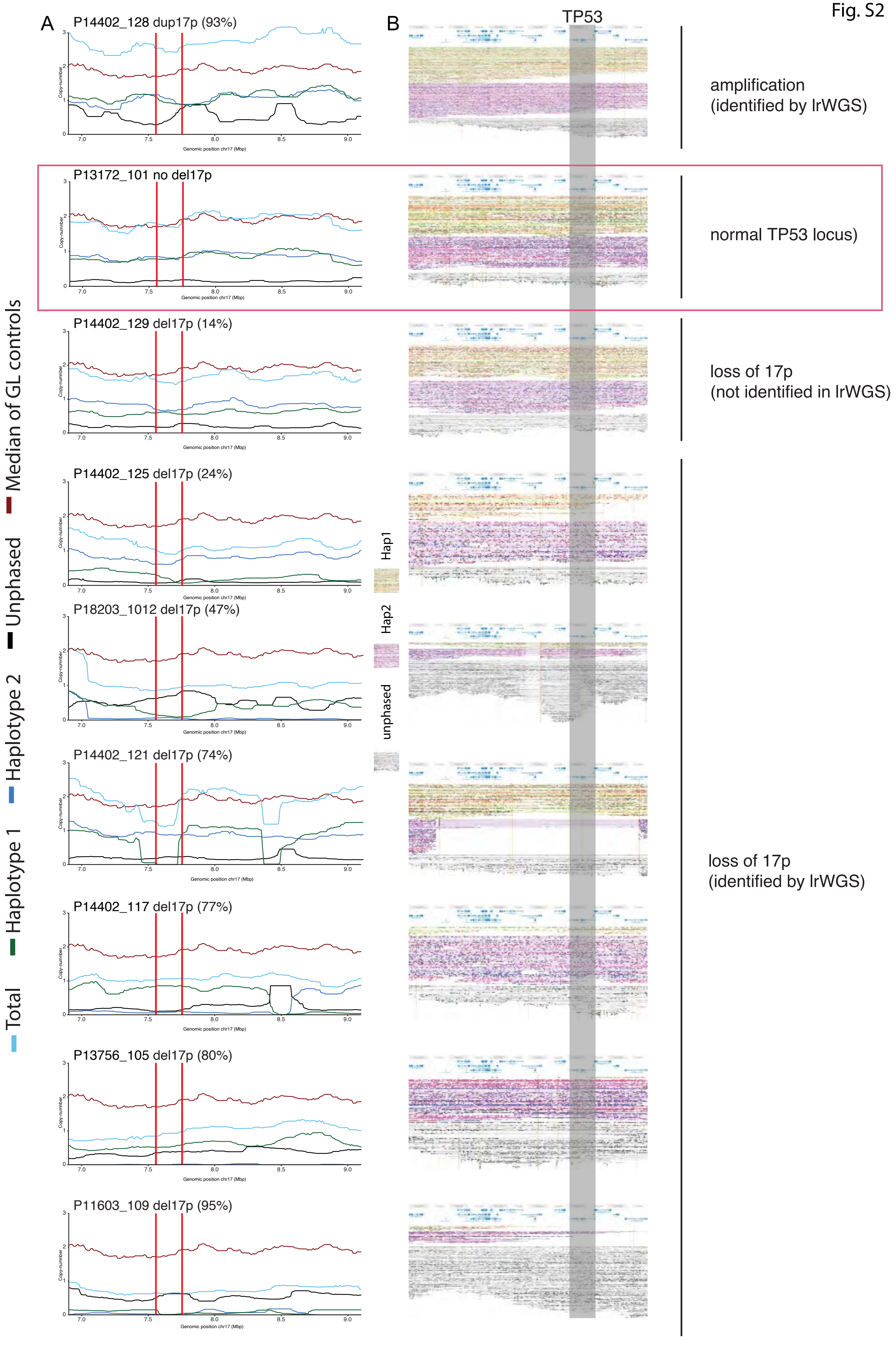
A
B
TP53
P14402_128 dup17p (93%)
P13172_101 no del17p
P14402_129 del17p (14%)
P14402_125 del17p (24%)
P18203_1012 del17p (47%)
P14402_121 del17p (74%)
P14402_117 del17p (77%)
P13756_105 del17p (80%)
P11603_109 del17p (95%)
Copy-number
Genomic position chr17 (Mbp)
Median of GL controls
Unphased
Haplotype 2
Haplotype 1
Total
Hap1
Hap2
unphased
amplification
(identified by lrWGS)
normal TP53 locus)
loss of 17p
(not identified in lrWGS)
loss of 17p
(identified by lrWGS)

**Figure S2. Linked-read WGS data allows for the identification of chr17p CNVs.** Copy-number (A) and phased read-clouds (B) are shown for eight patients with identified chr17p CNVs. In addition, one patient with a normal chr17p region is displayed for comparison. The percentage of cells with chr17p copy number aberration (as determined by FISH) is shown above the line graphs. (A) Line graph displaying the Copy-number of total, haplotype specific and unphased reads in the area surrounding the TP53 locus. Median copy-number of the normal control samples (GL, germline) is shown for comparison. Color coding is displayed at the left of the figure. Vertical lines indicate the boundaries of the FISH probes used to identify chr17p/TP53 CNVs. (B) Depiction of the phased read-clouds (drawn by Loupe). Read-clouds constituting haplotype 1 and haplotype 2 are indicated. The localization of the TP53 gene is indicated by the transparent grey box.

Fig. S3

A

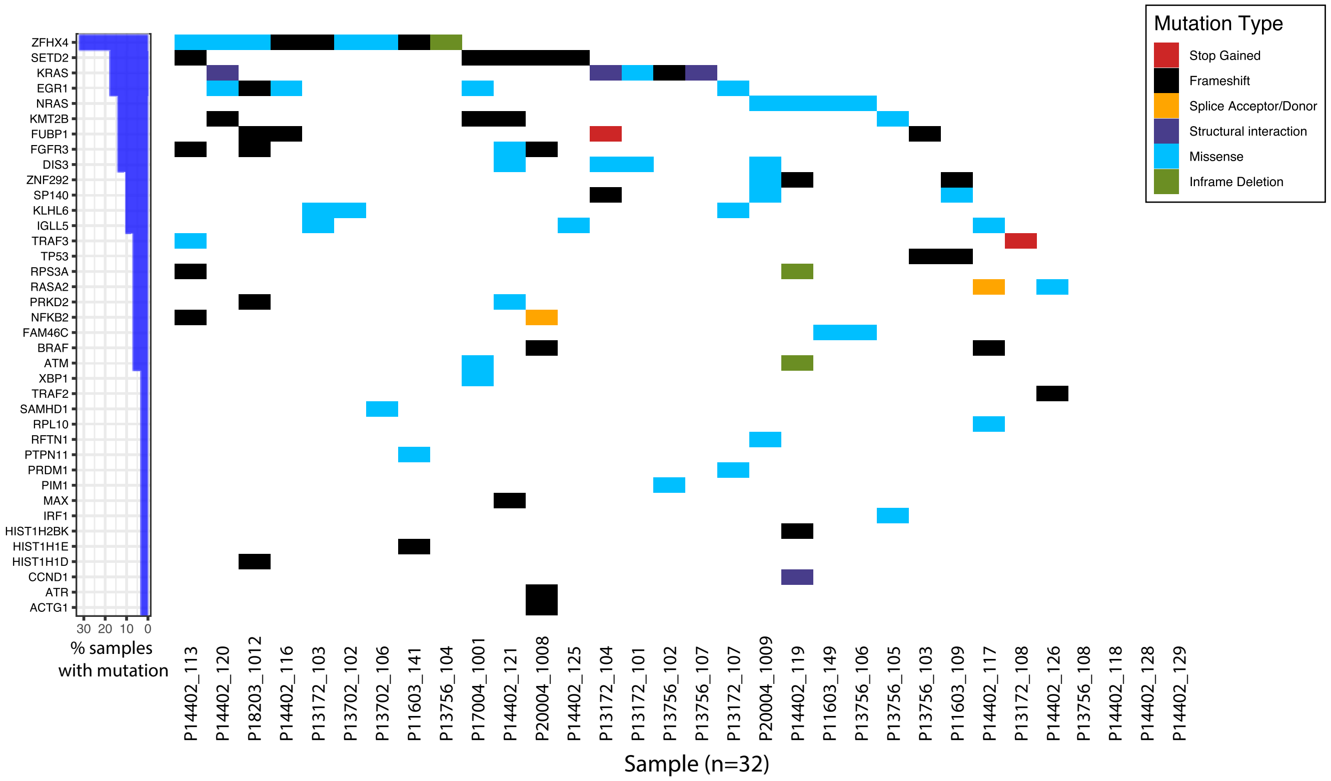

ZFHX4
SETD2
KRAS
EGR1
NRAS
KMT2B
FUBP1
FGFR3
DIS3
ZNF292
SP140
KLHL6
IGLL5
TRAF3
TP53
RPS3A
RASA2
PRKD2
NFKB2
FAM46C
BRAF
ATM
XBP1
TRAF2
SAMHD1
RPL10
RFTN1
PTPN11
PRDM1
PIM1
MAX
IRF1
HIST1H2BK
HIST1H1E
HIST1H1D
CCND1
ATR
ACTG1
30 20 10 0
% samples with mutation
P14402_113 P14402_120 P18203_1012 P14402_116 P13172_103 P13702_102 P13702_106 P11603_141 P13756_104 P17004_1001 P14402_121 P20004_1008 P14402_125 P13172_104 P13172_101 P13756_102 P13756_107 P13172_107 P20004_1009 P14402_119 P11603_149 P13756_106 P13756_105 P13756_103 P11603_109 P14402_117 P13172_108 P14402_126 P13756_108 P14402_118 P14402_128 P14402_129
Sample (n=32)
Mutation Type
Stop Gained
Frameshift
Splice Acceptor/Donor
Structural interaction
Missense
Inframe Deletion


B

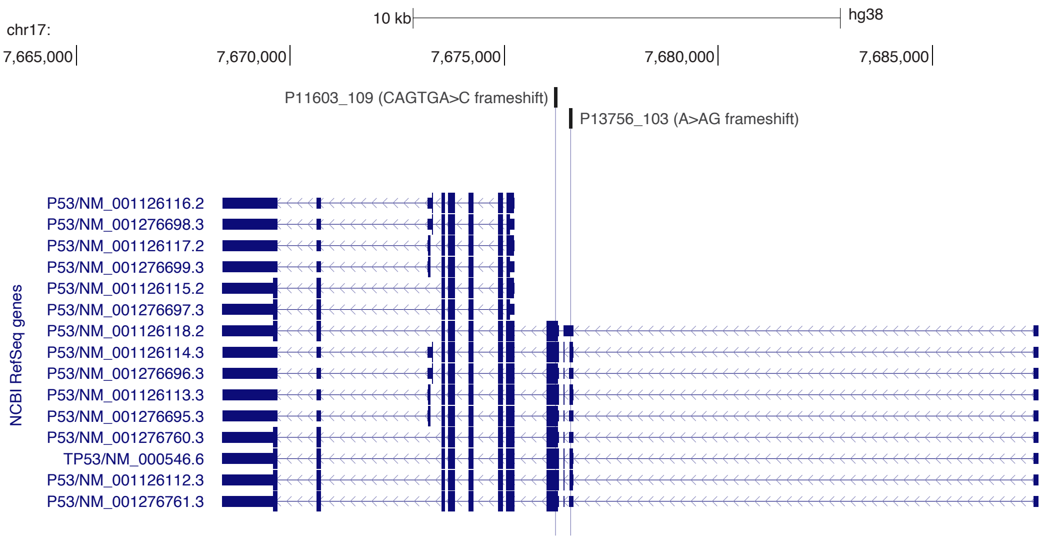

chr17:
10 kb
hg38
7,665,000
7,670,000
7,675,000
7,680,000
7,685,000
P11603_109 (CAGTGA>C frameshift)
P13756_103 (A>AG frameshift)
NCBI RefSeq genes
P53/NM_001126116.2
P53/NM_001276698.3
P53/NM_001126117.2
P53/NM_001276699.3
P53/NM_001126115.2
P53/NM_001276697.3
P53/NM_001126118.2
P53/NM_001126114.3
P53/NM_001276696.3
P53/NM_001126113.3
P53/NM_001276695.3
P53/NM_001276760.3
TP53/NM_000546.6
P53/NM_001126112.3
P53/NM_001276761.3

**Figure S3. Identification of common mutations and localization of the TP53 frame-shift mutations.** (A) Identified recurrent gene mutations in MM. (B) *TP53* sequence changes and localization in the exon structure of the TP53 gene.

Fig. S4

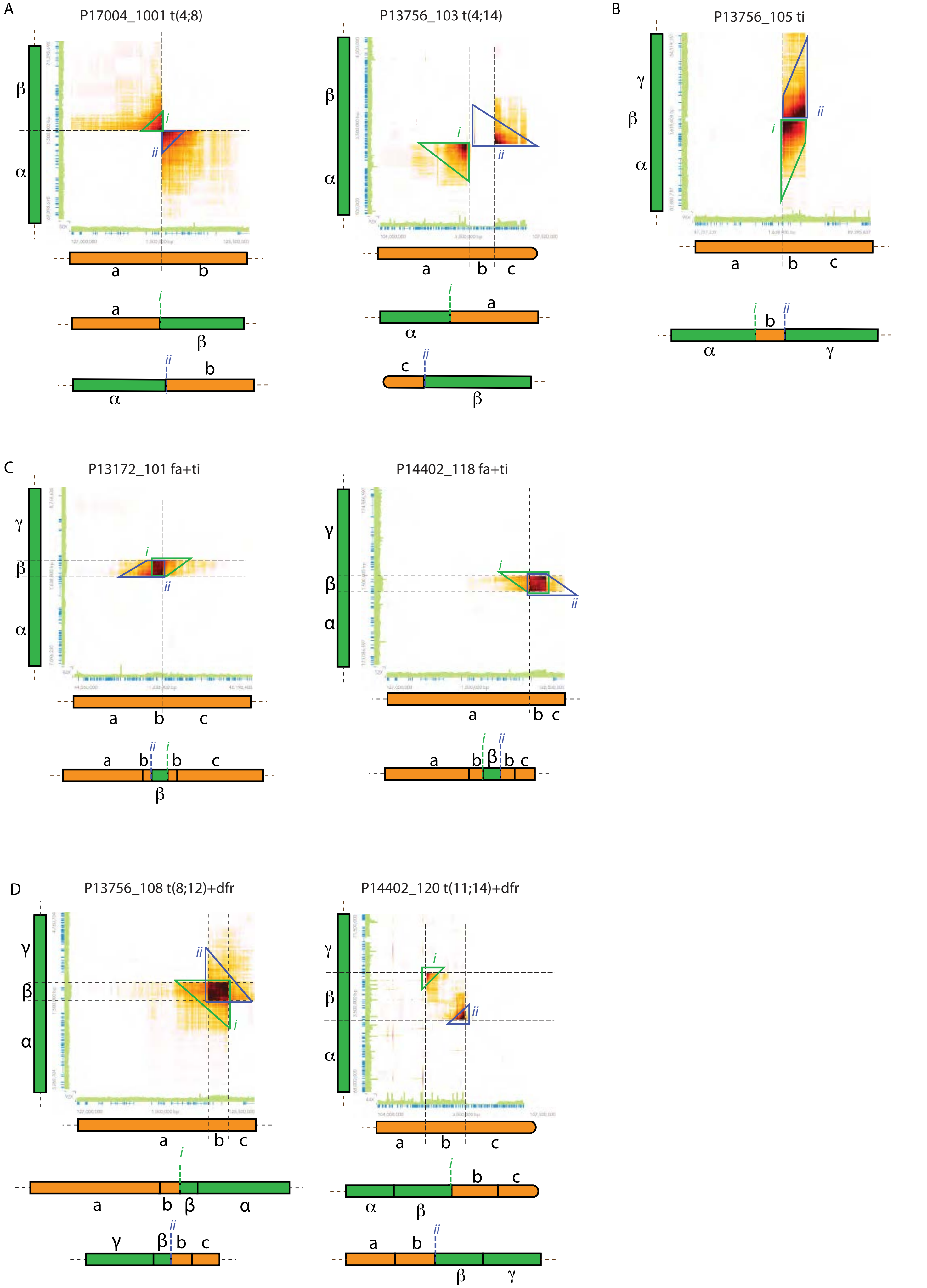
A
P17004_1001 t(4;8)
P13756_103 t(4;14)
B
P13756_105 ti
C
P13172_101 fa+ti
P14402_118 fa+ti
D
P13756_108 t(8;12)+dfr
P14402_120 t(11;14)+dfr

**Figure S4. Different structural variants generate distinct patterns in the lrWGS data.** (A-E) Heatmaps showing the barcode overlap for different structural variants (SVs) involving regions on different chromosomes and schematic representations of the SVs. Barcode overlap between indicated regions were drawn by Loupe with the yellow-red-black color scale indicating progressively higher barcode overlaps. As barcodes are essentially unique to each lrWGS read-cloud (and each read-cloud is generated from a single long DNA molecule), structural variants involving regions from different chromosomes give distinct patterns depending on the orientation and localization of the involved regions within the SV. Sequencing coverage (green bars) and the positions of coding regions (blue bars) is indicated on the axis of the heatmap. (A) Reciprocal translocations can be identified by clusters of read-clouds spanning the involved regions. This creates distinct triangular flair patterns 'pointing to' the translocation breakpoint (see green and blue triangles annotated as *i* and *ii* respectively). Orientation of the triangles depend on the orientation of the regions joined by the translocation (compare green and blue triangles in the examples in the left and right heatmap). The right example in addition to the translocation, harbors a deletion of the b region. Non-reciprocal translocations can be observed in a similar manner but present with only one 'triangle'. (B) Templated insertions (ti) create patterns with non-overlapping 'truncated triangles' as only a limited area (the inserted region) can share read-clouds with the surrounding chromosome. In the example, the insertion co-occurs with the loss of a small (the $\beta$ region) on the chromosome targeted by the insertion. The increased coverage of the inserted region can also be observed. (C) Templated insertions (ti) occurring in the middle of a focal amplification (fa), like simple insertions create 'truncated triangles' but these are in addition overlapping in the heatmap as the inserted region shares read-clouds with the fa region on either flank. Depending on the orientation of the insertion, the overlapping 'truncated triangles' will have different orientations (compare the left and right example). The increase in coverage caused by the fa and ti can be observed in both examples though it is more apparent in the case on the right. (D) Translocations with duplications of the flanking regions (dfr) can, similar to the fa with ti events (fa+ti), be observed as 'overlapping triangles'. However, as the regions are joined, the 'overlapping triangles' extend in both directions (i.e. truncated triangles are not created). Depending on the orientation of the translocation, the overlap of the 'triangles' is either visual (left) or only in terms of involved chromosomal regions (right). The increased coverage caused by dfr is most apparent in the case on the left.

Fig. S5

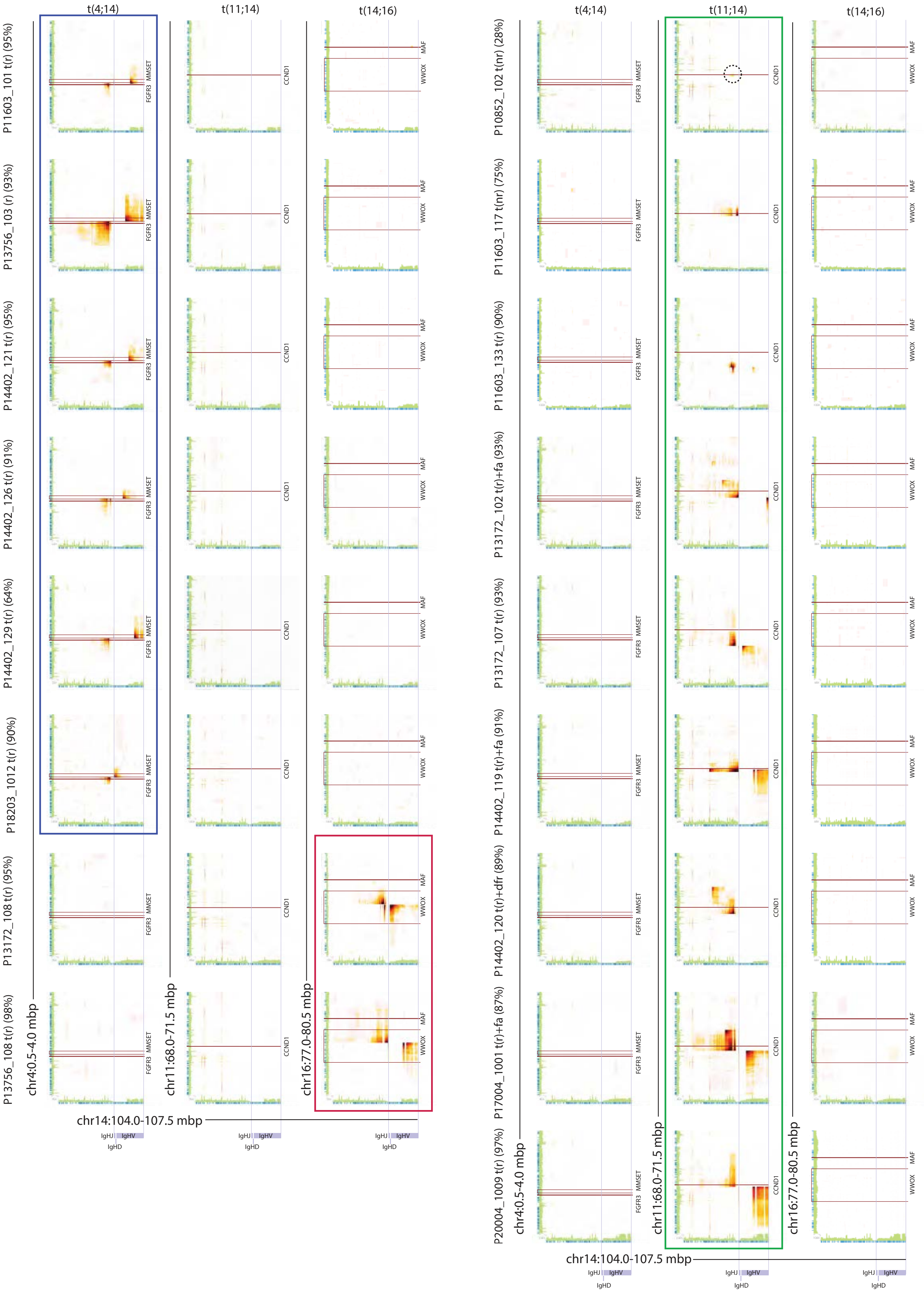

t(4;14)
t(11;14)
t(14;16)
P11603_101 t(r) (95%)
P13756_103 (r) (93%)
P14402_121 t(r) (95%)
P14402_126 t(r) (91%)
P14402_129 t(r) (64%)
P18203_1012 t(r) (90%)
P13172_108 t(r) (95%)
P13756_108 t(r) (98%)
P10852_102 t(nr) (28%)
P11603_117 t(nr) (75%)
P11603_133 t(r) (90%)
P13172_102 t(r)+fa (93%)
P13172_107 t(r) (93%)
P14402_119 t(r)+fa (91%)
P14402_120 t(r)+dfr (89%)
P17004_1001 t(r)+fa (87%)
P20004_1009 t(r) (97%)
FGFR3
MMSET
CCND1
MAF
WWOX
chr4:0.5-4.0 mbp
chr11:68.0-71.5 mbp
chr16:77.0-80.5 mbp
chr14:104.0-107.5 mbp
IgHJ
IgHV
IgHD

**Figure S5. Overview of identified t(4;14), t(11;14), and t(14;16) MM cases.** Heatmaps displaying the number of read-clouds shared between the IGH locus (chr14) and break-point regions on chr4 (MMSET), chr11 (CCND1), and chr16 (MAF) for all patients (n=17) with detected t(4;14), t(11;14), and t(14;16) translocations. Blue, red and green boxes indicate patients with verified t(4;14), t(11;14), and t(14;16) translocations respectively. Barcode overlap between indicated regions were drawn by Loupe with the yellow-red-black color scale indicating progressively higher barcode overlaps. Sequencing coverage (green bars) and the positions of coding regions (blue bars) is indicated on the axis of the heatmap. The type of translocation event is indicated next to the patient identifier: t(r), reciprocal translocation; t(nr), non-reciprocal translocation; t(r)+fa, reciprocal translocation with focal amplification; and t(r)+dfr, duplication of break-point flanking regions. The few read-clouds supporting the existence of the subclonal t(11;14) in P10852_102 are indicated by a dotted circle. Percentage (%) of cells found to have the indicated translocation by FISH is shown in parenthesis next to the patient identifier.

Fig. S6

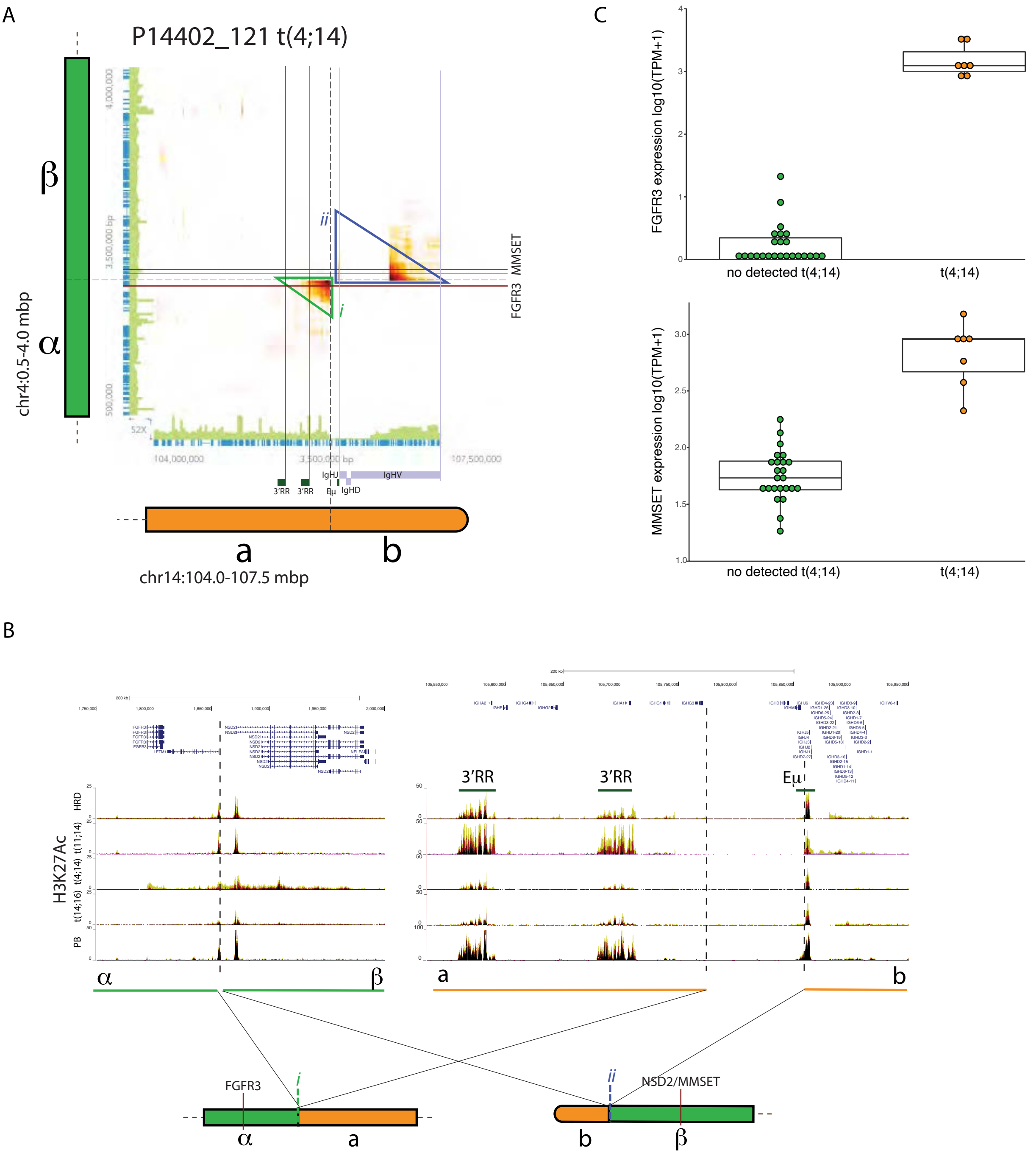
A
P14402_121 t(4;14)
β
α
chr4:0.5-4.0 mbp
ii
i
FGFR3
MMSET
3'RR
3'RR
Eμ
IgHJ
IgHD
IgHV
a
b
chr14:104.0-107.5 mbp
B
H3K27Ac
HRD
t(11;14)
t(4;14)
t(14;16)
PB
3'RR
3'RR
Eμ
α
β
a
b
FGFR3
i
ii
NSD2/MMSET
α
a
b
β
C
FGFR3 expression log10(TPM+1)
no detected t(4;14)
t(4;14)
MMSET expression log10(TPM+1)
no detected t(4;14)
t(4;14)

**Figure S6. Reciprocal t(4;14) translocations juxtapose IGH enhancers with the MMSET and FGFR3 loci.** (A) Heatmaps displaying the number of read-clouds shared between the IGH locus (chr14) and break-point regions on chr4 (MMSET) in P14402_121. Barcode overlap between indicated regions were drawn by Loupe with the yellow-red-black color scale indicating progressively higher barcode overlaps. Sequencing coverage (green bars) and the positions of coding regions (blue bars) is indicated on the axis of the heatmap. Position of the IGH VDJ regions and the 3'RR and Eμ enhancers are indicated. Green (*i*) and blue (*ii*) triangles indicate the position of the read-cloud clusters defining the structural event. (B) Overview of the enhancer landscape surrounding the translocation and schematic representation of derivate chromosomes. Genome browser tracks show the median H3K27Ac signal (red) ±1SD (yellow and black respectively) in normal plasma blasts (PB) and MM samples belonging to the indicated genetic group. (C) Expression of MMSET and FGFR3 in patients with and without t(4;14).

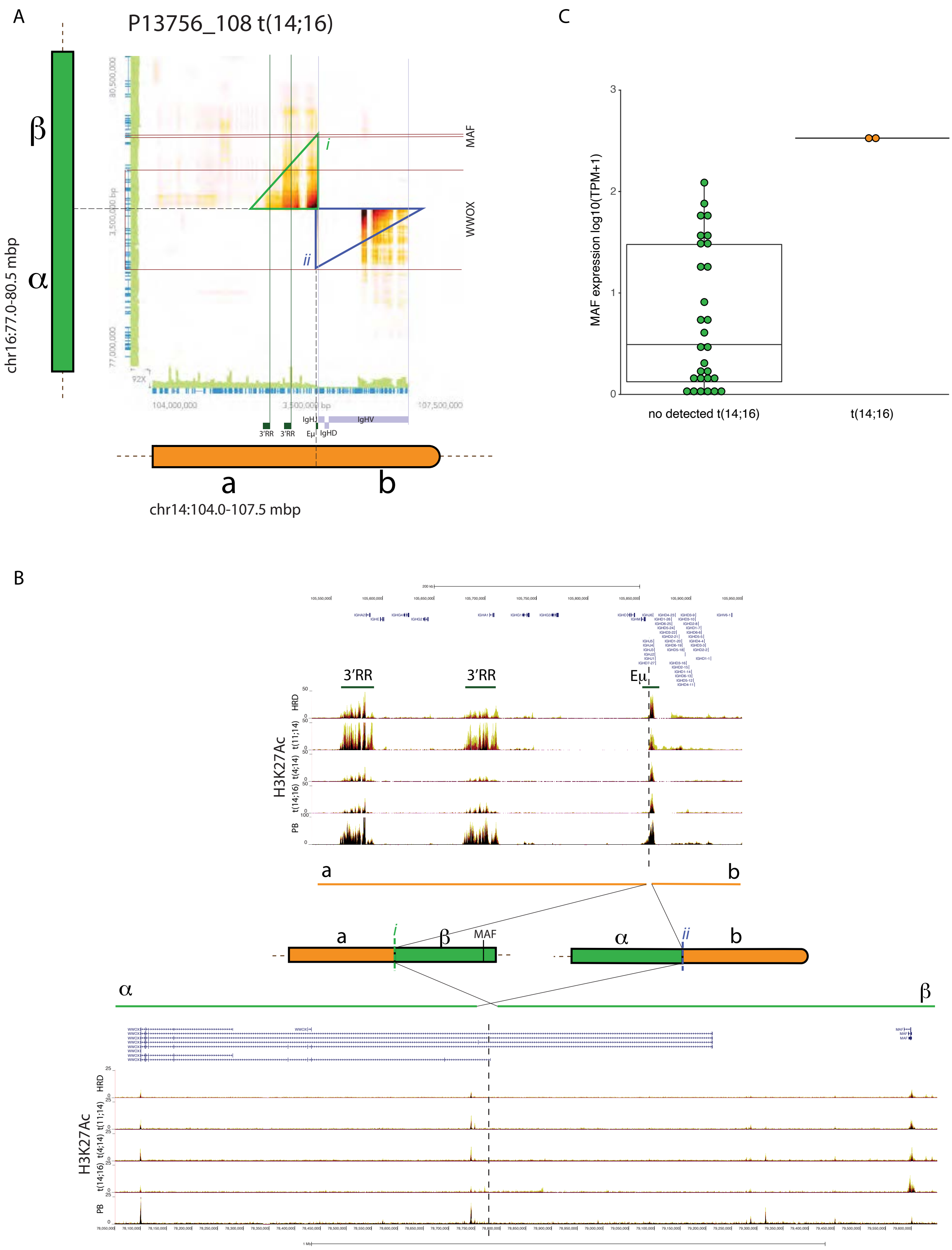
A
P13756_108 t(14;16)
β
α
chr16:77.0-80.5 mbp
MAF
WWOX
i
ii
3'RR
3'RR
IgHJ
IgHV
Eμ
IgHD
a
b
chr14:104.0-107.5 mbp
C
MAF expression log10(TPM+1)
no detected t(14;16)
t(14;16)
B
3'RR
3'RR
Eμ
H3K27Ac
HRD
t(11;14)
t(4;14)
t(14;16)
PB
a
b
a
β
MAF
α
b
α
β
HRD
t(11;14)
t(4;14)
t(14;16)
PB
H3K27Ac

**Figure S7. Reciprocal t(14;16) translocations juxtapose IGH enhancers with the MAF locus**. (A) Heatmaps displaying the number of read-clouds shared between the IGH locus (chr14) and break-point regions on chr16 (MAF) in P13756_108. Barcode overlap between indicated regions were drawn by Loupe with the yellow-red-black color scale indicating progressively higher barcode overlaps. Sequencing coverage (green bars) and the positions of coding regions (blue bars) is indicated on the axis of the heatmap. Position of the IGH VDJ regions and the 3'RR and Eμ enhancers are indicated. Green (*i*) and blue (*ii*) triangles indicate the position of the read-cloud clusters defining the structural event. (B) Overview of the enhancer landscape surrounding the translocation and schematic representation of derivate chromosomes. Genome browser tracks show the median H3K27Ac signal (red) ±1SD (yellow and black respectively) in normal plasma blasts (PB) and MM samples belonging to the indicated genetic group. (C) Expression of MAF in patients with and without t(14;16).

Fig. S8

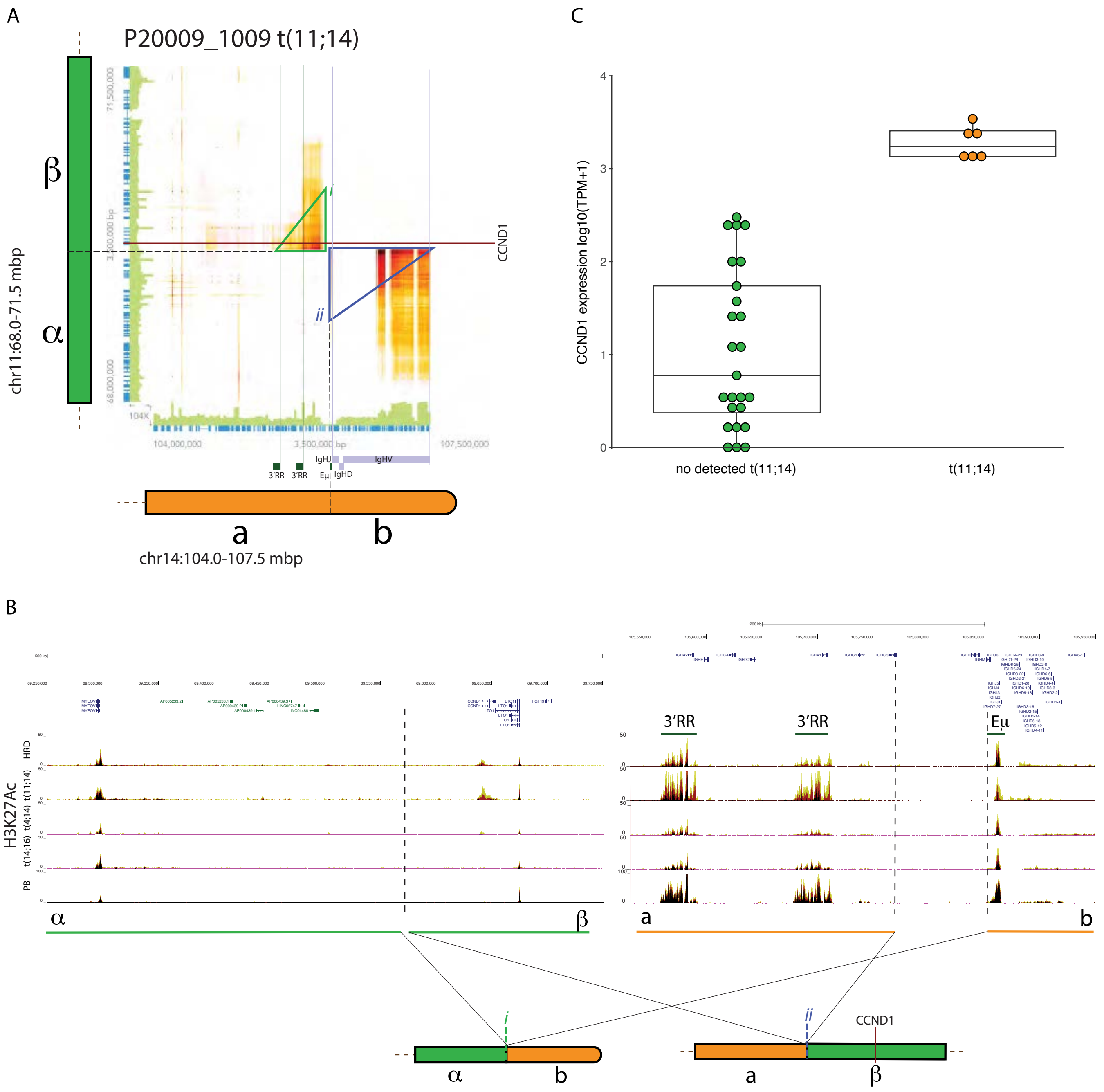

A
P20009_1009 t(11;14)
β
α
chr11:68.0-71.5 mbp
CCND1
i
ii
104,000,000
107,500,000
IgHJ
IgHV
3'RR
3'RR
Eμ
IgHD
a
b
chr14:104.0-107.5 mbp
B
H3K27Ac
HRD
t(11;14)
t(4;14)
t(14;16)
PB
3'RR
3'RR
Eμ
α
β
a
b
i
ii
CCND1
α
b
a
β
C
CCND1 expression log10(TPM+1)
0
1
2
3
4
no detected t(11;14)
t(11;14)

**Figure S8. Reciprocal t(11;14) translocations juxtapose IGH enhancers with the CCND1 locus**. (A) Heatmaps displaying the number of read-clouds shared between the IGH locus (chr14) and break-point regions on chr11 (CCND1) in P20009_1009. Barcode overlap between indicated regions were drawn by Loupe with the yellow-red-black color scale indicating progressively higher barcode overlaps. Sequencing coverage (green bars) and the positions of coding regions (blue bars) is indicated on the axis of the heatmap. Position of the IGH VDJ regions and the 3'RR and Eμ enhancers are indicated. Green (*i*) and blue (*ii*) triangles indicate the position of the read-cloud clusters defining the structural event. (B) Overview of the enhancer landscape surrounding the translocation and schematic representation of derivate chromosomes. Genome browser tracks show the median H3K27Ac signal (red) ±1SD (yellow and black respectively) in normal plasma blasts (PB) and MM samples belonging to the indicated genetic group. (C) Expression of CCND1 in patients with and without t(11;14).

### Fig. S9

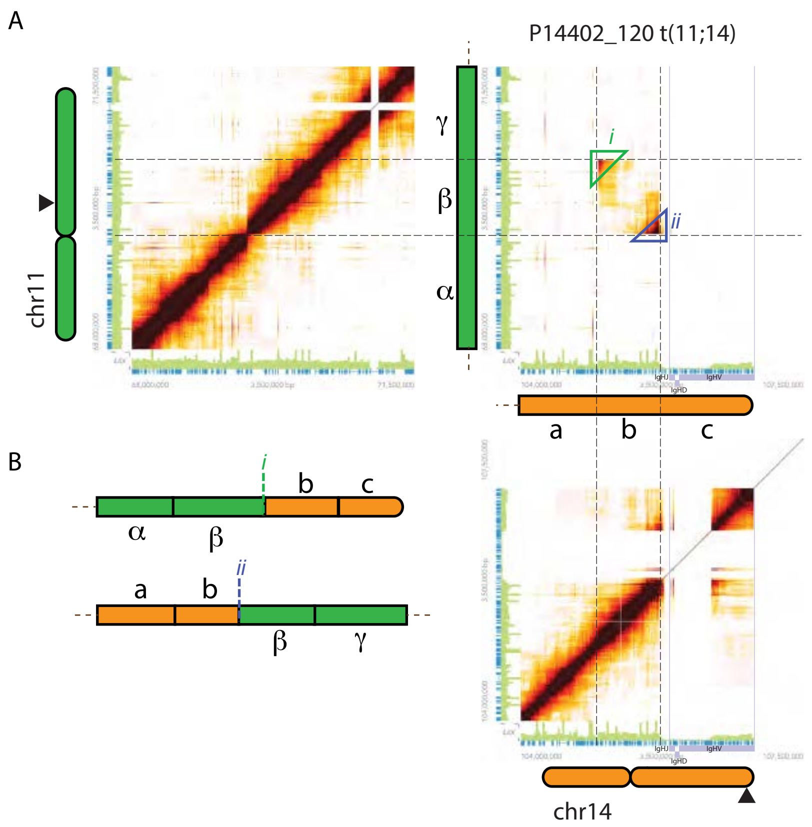

**Figure S9. The t(11;14) in P14402_120 carries duplications on both sides of the break-point.** (A) Heatmaps displaying the shared read-clouds on chr11 (top left), chr11 to chr14 (top right), and chr14 (bottom right) in P14402_120. Barcode overlap between indicated regions were drawn by Loupe with the yellow-red-black color scale indicating progressively higher barcode overlaps. Sequencing coverage (green bars) and the positions of coding regions (blue bars) is indicated on the axis of the heatmap. Triangles (*i*-*ii*) indicate the position of the read-cloud clusters defining the structural event. (B) Schematic representation of the derivative chromosomes generated by the t(11;14).

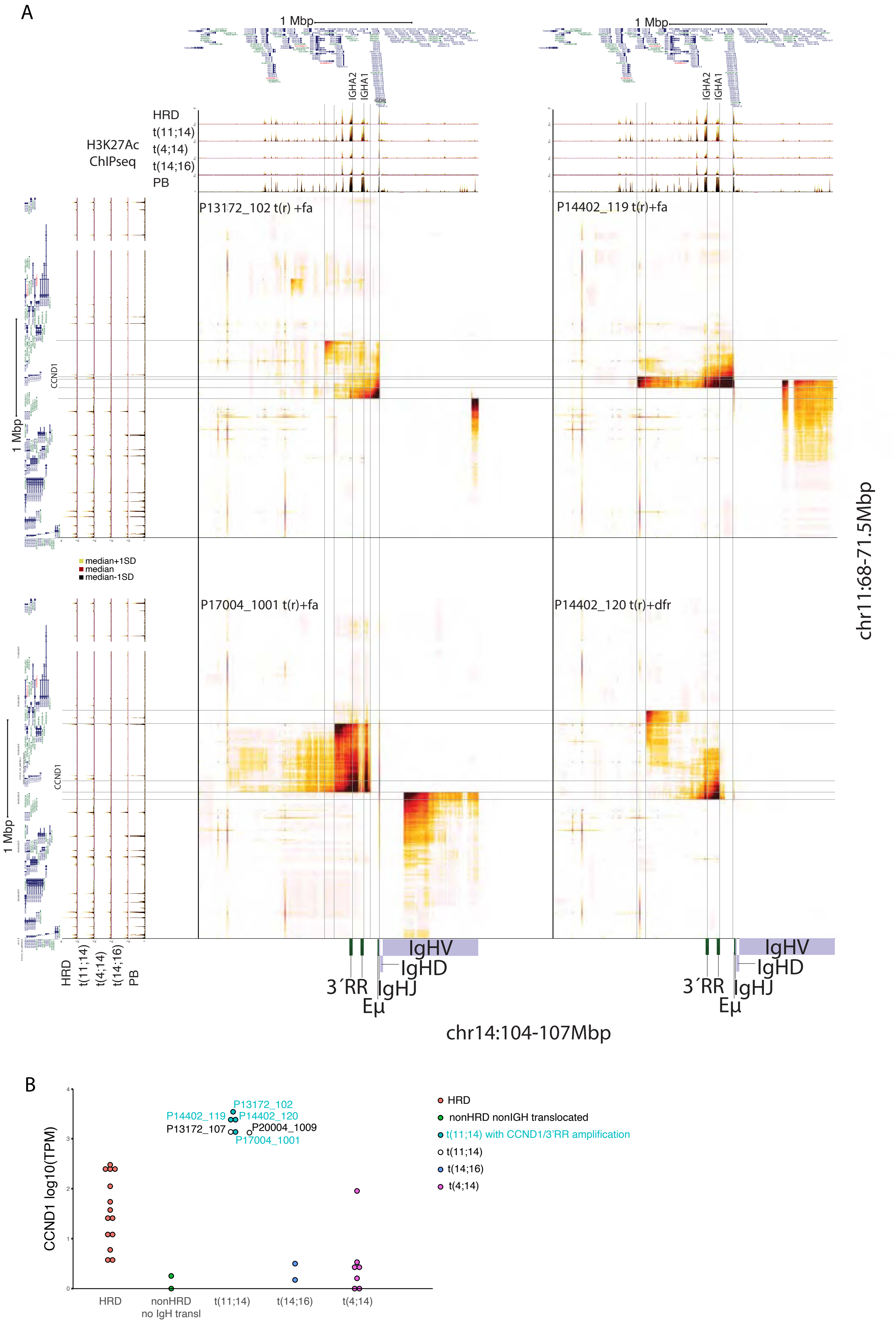
A
1 Mbp
IGHA2
IGHA1
HRD
t(11;14)
t(4;14)
t(14;16)
PB
H3K27Ac
ChIPseq
P13172_102 t(r) +fa
P14402_119 t(r)+fa
P17004_1001 t(r)+fa
P14402_120 t(r)+dfr
1 Mbp
CCND1
median+1SD
median
median-1SD
chr11:68-71.5Mbp
IgHV
IgHD
3´RR
IgHJ
Eμ
chr14:104-107Mbp
B
CCND1 log10(TPM)
P13172_102
P14402_119
P14402_120
P13172_107
P20004_1009
P17004_1001
HRD
nonHRD
no IgH transl
t(11;14)
t(14;16)
t(4;14)
HRD
nonHRD nonIGH translocated
t(11;14) with CCND1/3'RR amplification
t(11;14)
t(14;16)
t(4;14)

**Figure S10. The amplification of focal and flanking region in t(11;14) results in the duplication of the CCND1 and 3'RR enhancer regions.** (A) Heatmaps showing the number of shared read-clouds between chr11 and chr14 for the indicated patients. Barcode overlap between indicated regions were drawn by Loupe with the yellow-red-black color scale indicating progressively higher barcode overlaps. Tracks flanking the heatmaps (above and left) show the median H3K27Ac signal (red) $\pm$1SD (yellow and black respectively) in normal plasma blasts (PB) and MM samples belonging to the indicated genetic group. The boundaries of the breakpoints, the CCND1 locus and the IGHA-1/2 loci are indicated with light grey lines. The position of the IGH VDJ elements as well as the 3'RR and E$\mu$ enhancers (flanking the IGHA and IGHJ regions respectively) are indicated on chr14. (B) Expression of CCND1 in MM with the indicated genetic aberrations.

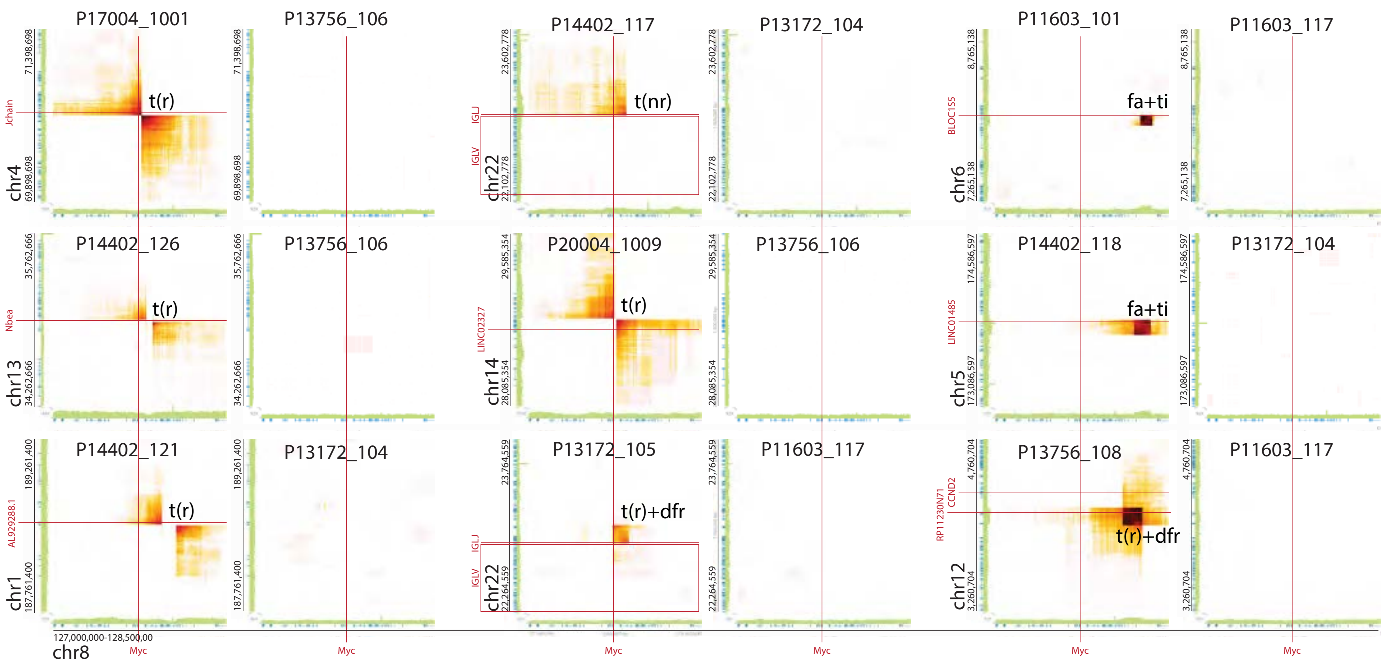
P17004_1001
P13756_106
P14402_117
P13172_104
P11603_101
P11603_117
P14402_126
P13756_106
P20004_1009
P13756_106
P14402_118
P13172_104
P14402_121
P13172_104
P13172_105
P11603_117
P13756_108
P11603_117
t(r)
t(nr)
fa+ti
t(r)+dfr
chr4
chr13
chr1
chr22
chr14
chr6
chr5
chr12
chr8
Jchain
Nbea
IGLJ
IGLV
LINC02327
BLOC1S5
LINC01485
CCND2
Myc
127,000,000-128,500,00

**Figure S11. The read-clouds supporting the existence of MYC SV are highly specific to the affected MMs.** Heatmaps displaying the number of read-clouds shared between the MYC locus and indicated genomic locations in MMs with (columns 1, 3 and 5) or without the detected SV (columns 2, 4 and 6). Barcode overlap between indicated regions were drawn by Loupe with the yellow-red-black color scale indicating progressively higher barcode overlaps. Sequencing coverage (green bars) and the positions of coding regions (blue bars) is indicated on the axis of the heatmap. The type of SV is indicated next to the read-cloud clusters supporting the existence of the SV: t(r), reciprocal translocation; t(nr), non-reciprocal translocation; t(r)+fa, reciprocal translocation with focal amplification; and t(r)+dfr, duplication of break-point flanking regions. Red vertical/horizontal lines show the position of indicated genes.

Fig. S12

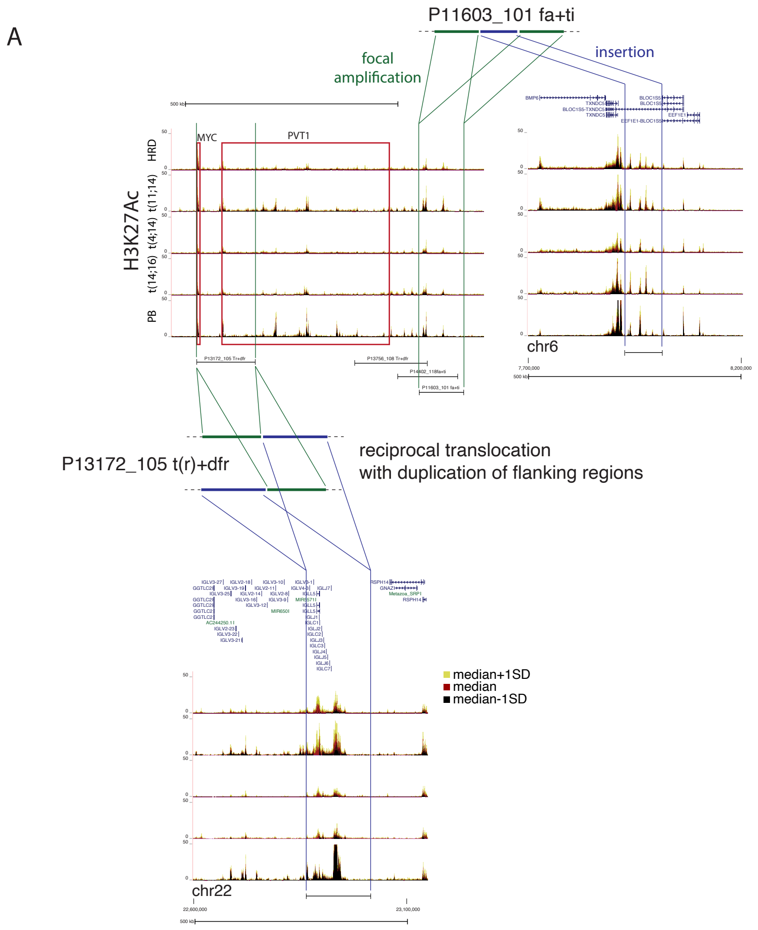
A
P11603_101 fa+ti
focal amplification
insertion
MYC
PVT1
H3K27Ac
HRD
t(11;14)
t(4;14)
t(14;16)
PB
chr6
P13172_105 t(r)+dfr
reciprocal translocation
with duplication of flanking regions
median+1SD
median
median-1SD
chr22

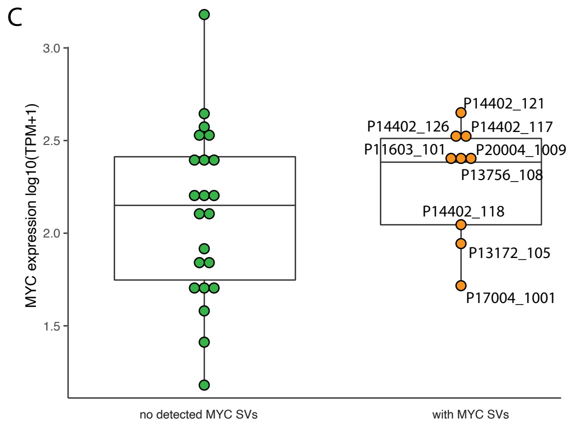
C
MYC expression log10(TPM+1)
3.0
2.5
2.0
1.5
P14402_121
P14402_126
P14402_117
P11603_101
P20004_1009
P13756_108
P14402_118
P13172_105
P17004_1001
no detected MYC SVs
with MYC SVs

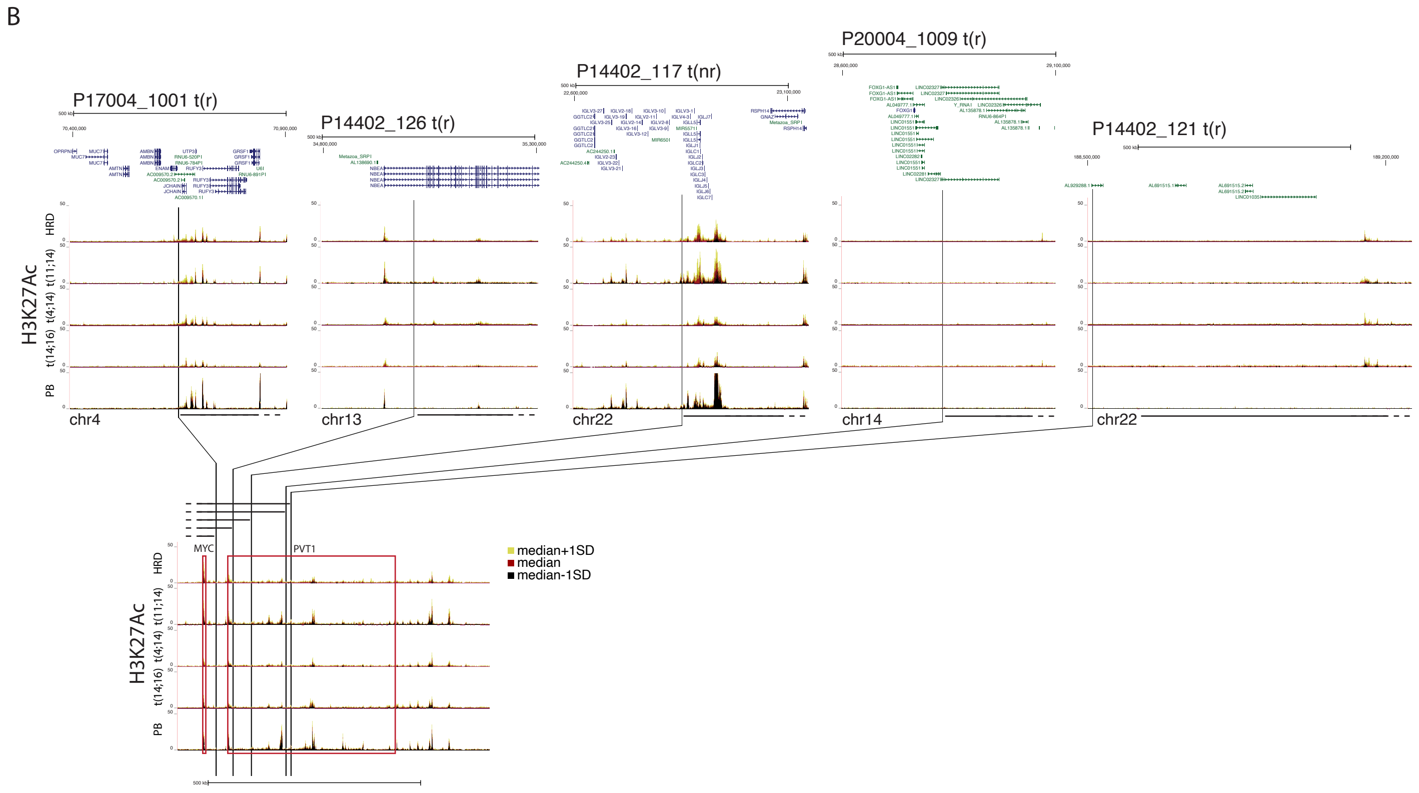
B
P17004_1001 t(r)
P14402_126 t(r)
P14402_117 t(nr)
P20004_1009 t(r)
P14402_121 t(r)
H3K27Ac
HRD
t(11;14)
t(4;14)
t(14;16)
PB
chr4
chr13
chr22
chr14
chr22
MYC
PVT1
median+1SD
median
median-1SD

**Figure S12. Overview of the genomic regions involved in the SVs effecting the MYC locus**. (A) Schematic representation of more complex SVs involving the MYC locus and tracks showing the median H3K27Ac signal (red) ±1SD (yellow and black respectively) in the involved regions in indicated types of MM and PBs. (B) Representation of the regions involved in simple translocations (reciprocal or non-reciprocal) affecting the MYC locus and tracks showing the H3K27Ac signals (median±SD) in the involved regions in indicated types of MM and PBs. The regions joined on the derivate chromosome containing the MYC locus are indicated by black lines. The type of SV effecting the individual MM is indicated next to the patient identifier: t(r), reciprocal translocation; t(nr), non-reciprocal translocation; t(r)+fa, reciprocal translocation with focal amplification; and t(r)+dfr, duplication of break-point flanking regions. (C) Expression of MYC in patients with and without SVs involving the MYC locus.

Fig. S13

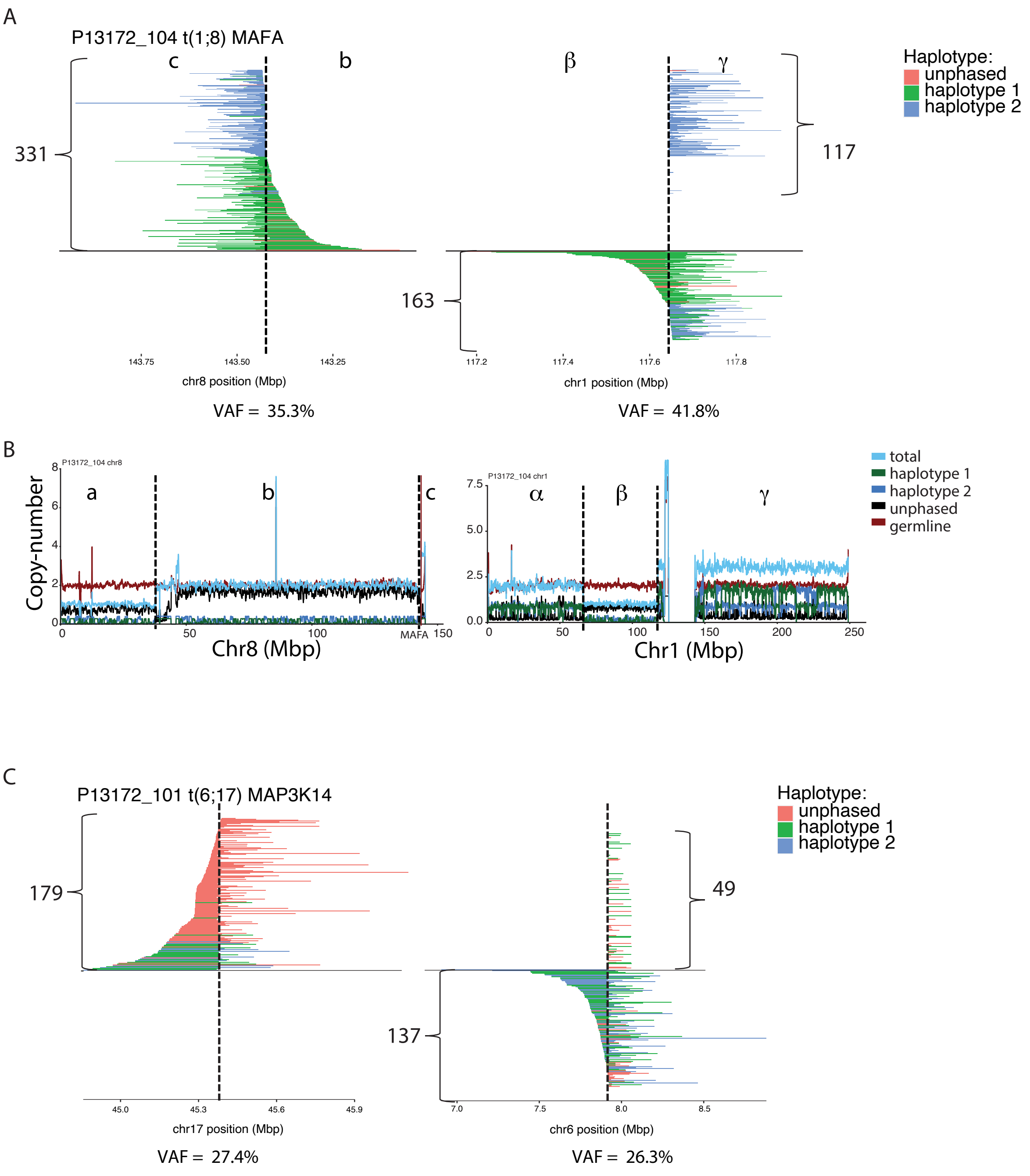
A
P13172_104 t(1;8) MAFA
c
b
331
β
γ
117
163
Haplotype:
unphased
haplotype 1
haplotype 2
143.75
143.50
143.25
chr8 position (Mbp)
117.2
117.4
117.6
117.8
chr1 position (Mbp)
VAF = 35.3%
VAF = 41.8%
B
P13172_104 chr8
Copy-number
a
b
c
MAFA
Chr8 (Mbp)
P13172_104 chr1
α
β
γ
Chr1 (Mbp)
total
haplotype 1
haplotype 2
unphased
germline
C
P13172_101 t(6;17) MAP3K14
179
49
137
Haplotype:
unphased
haplotype 1
haplotype 2
45.0
45.3
45.6
45.9
chr17 position (Mbp)
7.0
7.5
8.0
8.5
chr6 position (Mbp)
VAF = 27.4%
VAF = 26.3%

**Figure S13. Analysis of the read-clouds in the t(1;8) MAFA and t(6;17) MAP3K14 break-point regions.** (A) Localization of read-clouds involving the breakpoint regions of the t(1;8) involving the MAFA locus in MM P13172_104. VAF at either break-point site (indicated by black dotted lines) were calculated by dividing the number of read-clouds spanning chr1 to chr8 with the total number of read-clouds involving $\pm$10kb of the breakpoint. (B) Copy-number of chr8 and chr1 on patient P13172_104. Considering the amplification of the chr8 c and chr1 $\gamma$ regions (with 3 copies or more), the VAF of $\gamma$ to c spanning read-clouds suggest that the SV is clonal. (C) Localization of the read-clouds involving the breakpoint regions constituting the t(6;17) near the MAP3K14 locus in MM P13172_101. VAF at either break-point site (indicated by black dotted lines) were calculated by dividing the number of read-clouds spanning chr6 to chr17 with the total number of read-clouds involving $\pm$10kb of the breakpoint. Considering the normal copy-number of both involved chromosomes (data not shown), the VAF calculated at both breakpoints suggests that the SV is subclonal.
